# Limb, joint and pelvic kinematic control in the quail coping with step perturbations

**DOI:** 10.1101/2022.01.11.475813

**Authors:** Emanuel Andrada, Oliver Mothes, Heiko Stark, Matthew C. Tresch, Joachim Denzler, Martin S. Fischer, Reinhard Blickhan

## Abstract

Small cursorial birds display remarkable walking skills and can negotiate complex and unstructured terrains with ease. The neuromechanical control strategies necessary to adapt to these challenging terrains are still not well understood. Here, we analyzed the 2D- and 3D pelvic and leg kinematic strategies employed by the common quail to negotiate visible step-up and step-down perturbations of 1 cm, 2.5 cm, and 5 cm. We used biplanar fluoroscopy to accurately describe joint positions in three dimensions and performed semi-automatic landmark localization using deep learning.

Quails negotiated vertical perturbations without major problems and rapidly regained steady-state locomotion. When coping with step-up perturbations, the quail mostly adapted the trailing limb to permit the leading leg to step on the elevated substrate in a similar way as it did during level locomotion. When the quail negotiated step-down perturbations, both legs showed significant adaptations. For small and moderate perturbations (not inducing aerial running) the quail kept the function of the distal joints (i.e., their kinematic pattern) largely unchanged during uneven locomotion, and most changes occurred in proximal joints. The hip regulated leg length, while the distal joints maintained the spring-damped limb patterns. However, to negotiate the largest visible step perturbations, more dramatic kinematic alterations were observed. For these large perturbations, all joints contributed to leg lengthening/ shortening in the trailing leg and both the trailing and leading legs stepped more vertically and less abducted. This indicates a shift from a dynamic walking program to strategies that are focused on maximizing safety.

## Introduction

Encompassing almost ten thousand species, birds (clade Aves) are the most successful bipeds. Despite their flying abilities, they also represent a valuable study group to understand adaptations to terrestrial locomotion. For example, there are bird species that combine remarkable flying and walking abilities (e.g., waders ^1,2^). Other species evolved to live on the ground, losing partially or completely their ability to fly. Within the latter group encompassing about sixty species, the quail (*Coturnix coturnix*), is representative for the group of small cursorial birds. Like most of this group, the quail prefer grounded running (a running gait without aerial phases) during unrestricted level locomotion ^3,4^ In the wild, however, the quail must navigate over complex and unstructured terrains. Locomotion might become non-periodic, altering the kinematic and mechanical demands placed on the neuromechanical control system as compared to level locomotion. Our understanding of how animals’ neuromechanical control strategies adapt to these changing demands, despite important progress achieved in the past years, remains elusive.

It is believed that animals combine the intrinsic stability of their body mechanics with their neuronal control to negotiate rough terrains. The assumption is that anticipatory (feedforward) mechanisms pre-adjust limb kinematics and impedance before the leg contacts the ground, to reduce the need for reactive (feedback) response to readapt posture during stance ^5–9^. In the last years, two dimensional neuromechanical studies have tried to bring light to the adaptive mechanisms underlying uneven locomotion in the bird. Results of those studies showed that birds use anticipatory maneuvers to vault upwards in order to avoid excessive crouched postures on an obstacle ^10,11^. Birds also use leg retraction in late swing to regulate landing conditions ^10,12^, to minimize fluctuations in leg loading during uneven locomotion ^13^, and to prevent falls ^14,15^. Late-swing retraction is known to increase stability of locomotion as it changes the angle of attack of the leg at touch down (TD) according to obstacle height ^16^. In small birds, the retraction of the leading leg can be the consequence of the leg placement strategy called fixed aperture angle ^4^. In this strategy, the angle between the leg going to contact on the ground (usually termed leading) and the supporting legs (usually termed trailing) is fixed before TD. The retraction of the leading leg is thus automatically adapted for locomotion speed ^4,17,18^. The aperture angle strategy has not yet been tested in birds facing perturbations, although there is some evidence for its use by humans during uneven locomotion^19^.

Interestingly, the guinea fowl (*Numida meleagris*) did not exhibit anticipatory strategies for negotiating obstacles on a treadmill ^9,20^. This result indicates a robust inherent stability that was also shown in the ability of birds to cope with camouflaged drops ^12^. The robustness of avian level locomotion was assessed using a simple model including an effective leg (the segment spanning from the hip to the toe, Fig. 1F) and a trunk ^18^. The model produced self-stable gaits and was able to cope with steps over obstacles or sudden drops without the need for feedback control or even the need for tuning feedforward strategies ^18,21^.

**Figure 1.**
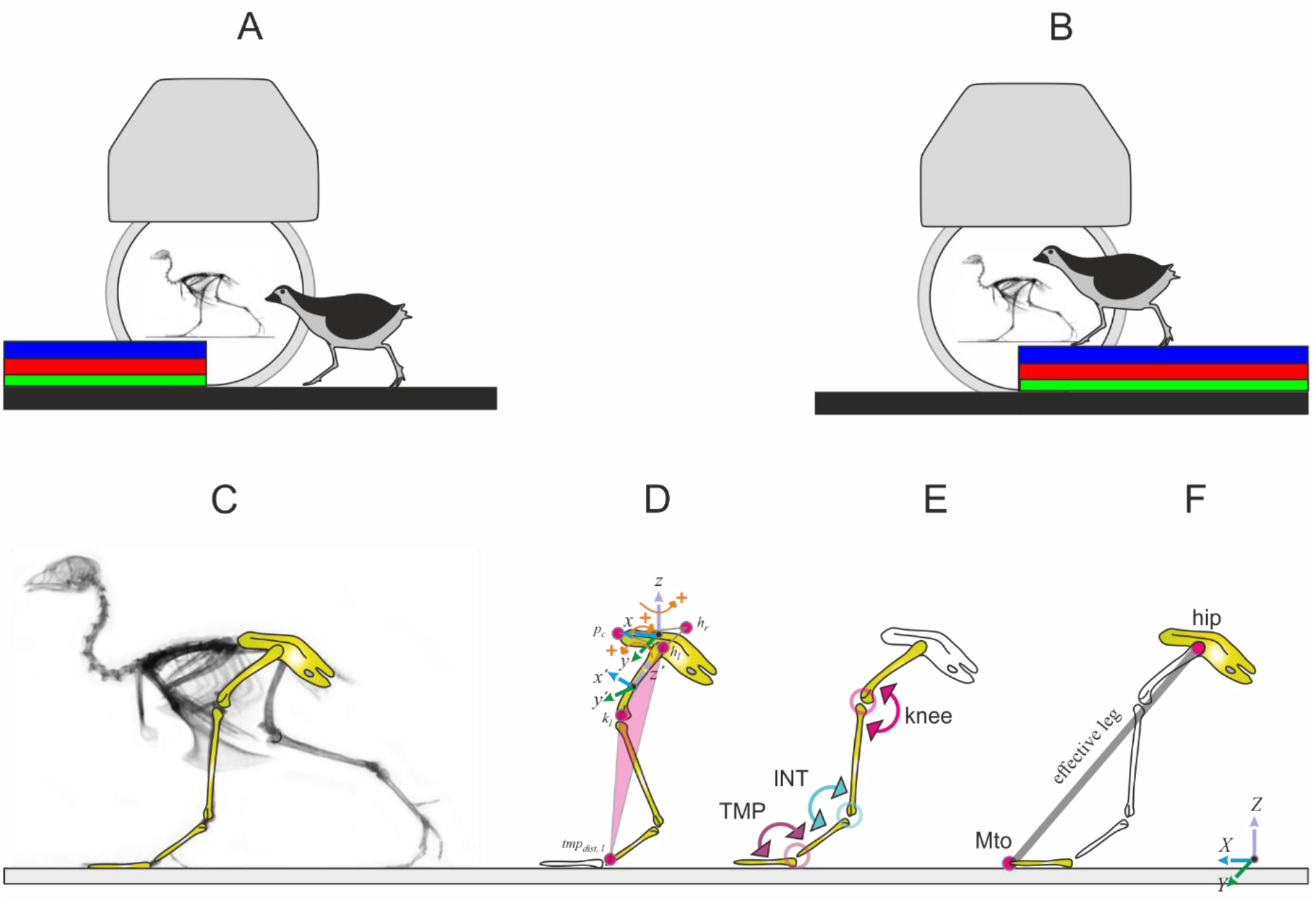
Experimental setup and 2D / 3D global and joint limbs kinematics. The quail negotiated visible step-up (A) and step down (B) perturbations of 1 cm (green), 2.5 cm (red), and 5 cm (blue) height. Body and hindlimb kinematics were captured using biplanar fluoroscopy. C) analyzed body segments. D) 3D kinematics of the pelvis relative to the global coordinate system, and rotation of the whole leg related to the pelvis. The last estimates the three-dimensional rotations occurring at the hip joint. The whole leg is a plane formed by the hip (e.g., h_l_), the knee (e.g., k_l_) and the distal marker of the tarsometatarsus (tmt_dist.l_), see methods, E) joint kinematics (INT: intertarsal joint, TMP: tarsometatarsal-phalangeal joint), F) effective leg (Mto: tip of the middle toe).

To our knowledge, there is no previous literature on three-dimensional analyses of avian locomotion over uneven surfaces. Even for level locomotion, three-dimensional analyses of avian locomotion are uncommon e.g., ^22–25^.

In this study, we aimed to uncover pelvic, leg, and joint kinematic adaptations to visible vertical perturbations (step up and step down, Fig. 1), and how these adaptations influence leg response after TD. We searched for relationships between simple model representations of the leg and joint kinematics.

Simple model representations like the effective leg help to understand basic strategies for stability or economy of locomotion e.g., ^4,5,17,26–28^ and can be used as global goals for the control of limb joints ^29^. During unrestricted locomotion there is evidence of an interplay between effective leg and limb segmental angles. In humans, Japanese macaques and the quail, limb segmental angles (thigh, shank, and foot) covary in a way that they form a planar loop in a three-dimensional space ^30–34^. This result indicates that intersegmental coordination might reduce the number of degrees of freedom to control the leg from three (i.e., joint angles) to two (i.e., effective leg length and angle).

Due to the redundant nature of the segmented leg, different combinations of joint kinematics can lead to the same effective leg length and angle before TD, but to differing leg responses later during stance. Thus, we can expect that their combined analysis helps to infer quail motor control goals on rough terrains. In our experiments, we used biplanar fluoroscopy to accurately describe joint positions in three dimensions (Fig. 1 A, B). Because of our constrained field of view, we focused our analysis on preadaptation strategies, i.e., from the stride i-1 (before perturbation) to stride i (in perturbation).

We expected perturbation-type (up vs. down) and perturbation-height related changes in leg kinematics, as animals preadapt and redirect the body when negotiating a visible vertical perturbation. While kinematics cannot predict dynamics, we anticipated that the knowledge of the interaction between kinematics and dynamics during level locomotion could help us to deduce joint related pre/post adaptations and thus to infer the main goals of neuromechanical strategies used by animals to cope with visual vertical perturbations.

Our main predictions were the following: 1) the effective leg kinematics will be unchanged for small perturbations, 2) these adaptations will be made through adjustments primarily in proximal joints, and 3) for larger perturbations that compromise safety, distinct adaptations might be required in both leg and joint levels.

## Results

Quails negotiated vertical perturbations ranging from ca. 10% to 50% of their effective leg length without major problems. None of the subjects lost visible stability or stumbled because of the perturbations. Furthermore, they recovered from perturbations after one or two steps. To overcome 1 cm vertical perturbations quails usually switched to aerial running for both step-up and step-down perturbations. For negotiating 2.5 cm and 5 cm perturbations quails relied on double support phases, except for 5 cm drops, where they switched sometimes to aerial running after the perturbation. On average, locomotion speed decreased, while contact and swing times tended to increase with perturbation height (Table 1), although during step-up locomotion, contact and swing times for 2.5 cm height were longer than those measured for 5 cm height.

**Table 1.** spatiotemporal parameters

|  |  | Step up |  |  | Step down |  |  | Level |
| --- | --- | --- | --- | --- | --- | --- | --- | --- |
|  |  | 1 cm | 2.5 cm | 5 cm | 1 cm | 2.5 cm | 5 cm |  |
| speed [m s<sup>-1</sup>] |  | 0.65 ±0.12 | 0.55 ±0.2 | 0.51 ±0.16 | 0.94 ±0.27 | 0.51 ±0.24 | 0.44 ±0.17 | 0.6±0.11 |
| Contact time [s] | trailing | 0.23 ±0.03 | 0.30 ±0.12 | 0.25 ±0.06 | 0.18 ±0.04 | 0.25 ±0.18 | 0.34 ±0.09 | 0.22 ± 0.05 |
|  | leading | 0.22 ±0.03 | 0.33 ±0.19 | 0.29 ±0.06 | 0.15 ±0.04 | 0.21 ±0.06 | 0.21 ±0.06 |  |
| Swing time [s] | trailing | 0.17 ±0.1 | 0.23 ±0.12 | 0.20 ±0.03 | 0.14 ±0.01 | 0.19 ±0.03 | 0.14 ±0.03 | 0.14± 0.04 |
|  | leading | 0.17 ±0.1 | 0.22 ±0.1 | 0.17 ±0.04 | 0.17 ±0.01 | 0.20 ±0.02 | 0.20 ±0.05 |  |

In the following only selected significant differences are presented, please refer to the tables for further information about significance values.

### Analysis of effective leg kinematics

#### Stepping up, trailing leg

Overall patterns of the effective leg length for the trailing limb were similar for level and step-up locomotion. After TD, the supporting effective leg is compressed, then slightly extended until toe-off (TO). During the swing, the leg shortened and rapidly extended until the next TD. However, some differences can be observed between level and step-up locomotion. Quails prepare step-up TD with longer effective trailing legs than observed during level locomotion. During stance, step-up perturbations increased trailing leg extension and reduced leg retraction significantly (see Fig. 2 and Table 2).

**Figure 2.**
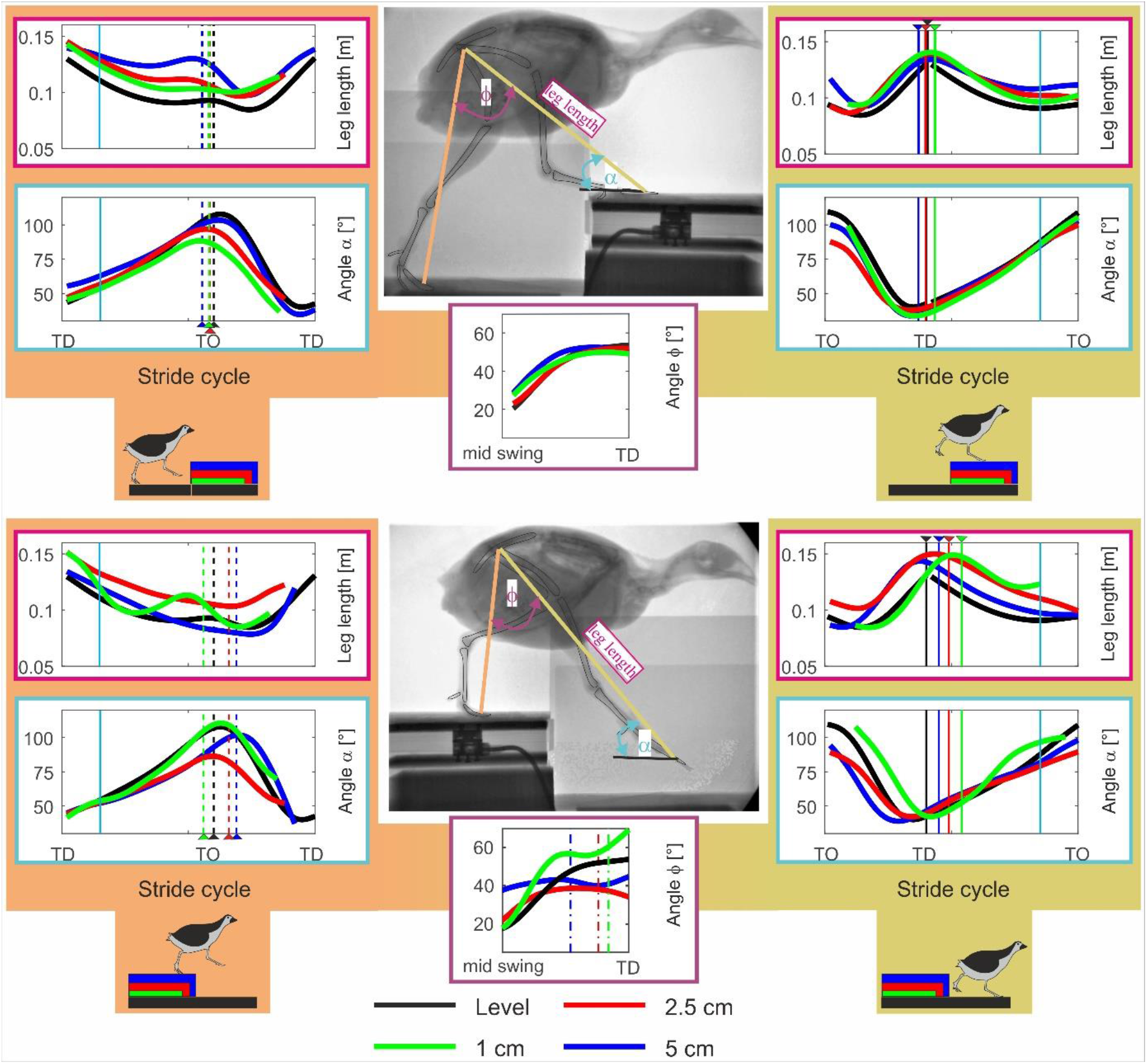
Effective leg kinematics. Effective leg length, effective leg angle and aperture angle between effective legs. level (black) and step locomotion (1 cm: green, 2.5 cm: red, 5 cm: blue) in the quail. Rows 1 and 4 display the effective leg length. Rows 2 and 5 effective leg angle (α). Single figures in Rows 3 and 6 display the aperture angle φ. Left: trailing leg stepping before the vertical perturbation (i-1), right: leading leg stepping after the vertical perturbation (i). Curves display mean values. Black, blue, red, green dashed lines indicate toe-off (TO), while solid lines touch down (TD). Cyan solid lines indicate 15% and 85% of the stride.

**Table 2.**
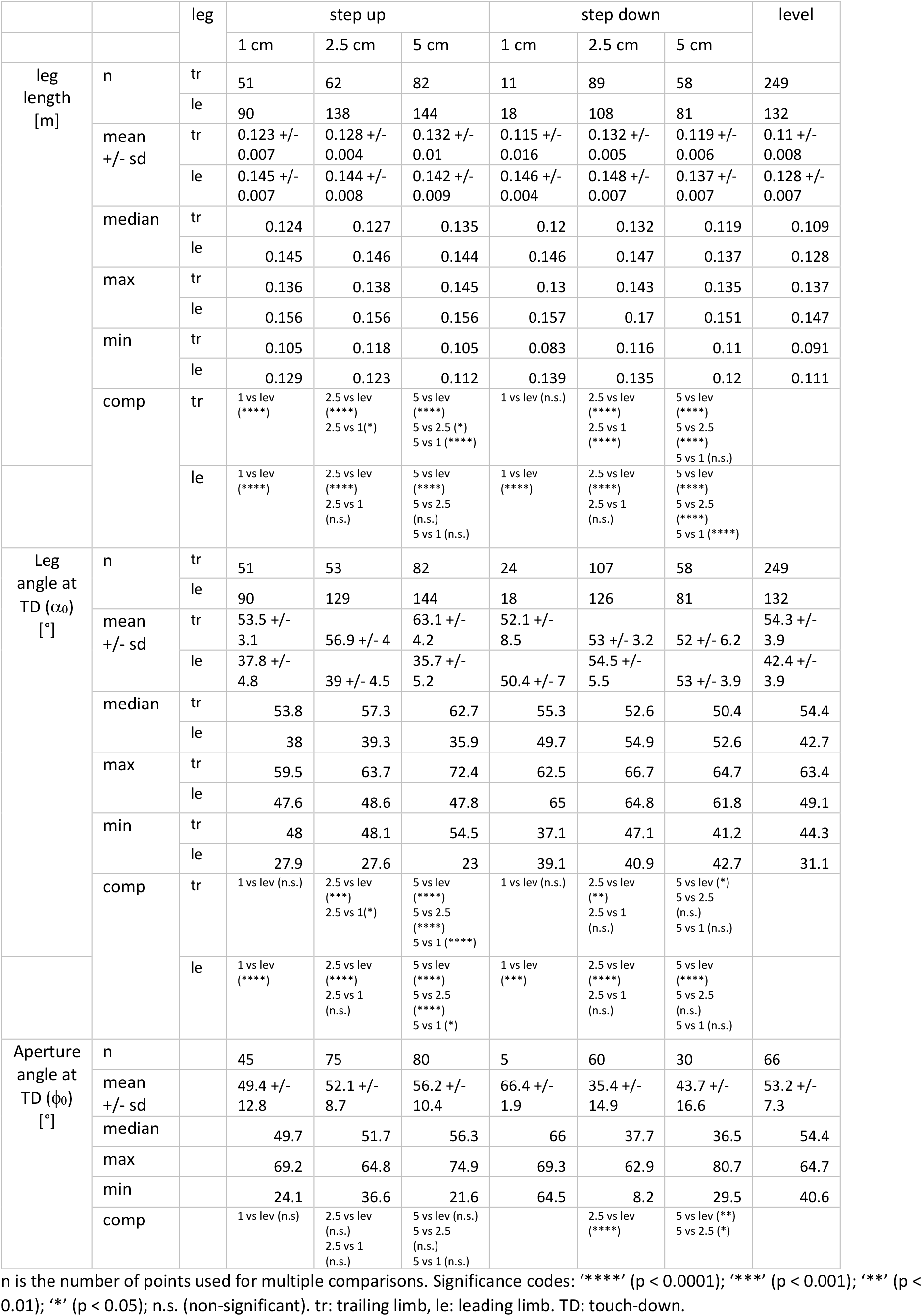
Mean, median, max, min values and multiple comparisons for the effective leg during level and step locomotion. For the trailing limb, analyses were performed at early stance (15% of the stride ± 4%). For the leading limb, around TD (TD ± 4%).

#### Stepping up, leading leg

In general, the effective kinematics of the leading leg during step-up locomotion were similar to those observed during level locomotion. No significant adaptations in the leading leg can be observed in the effective leg length before and at TD on the step, although after mid-swing the effective leg length is slightly longer during step-up locomotion as compared to level locomotion. Although the trajectory of the effective leg angle on the step was not substantially altered as compared to level locomotion, some minor differences can be observed. For example, the leading leg starts the swing phase more vertically oriented and contacts the elevated substrate with a slightly less vertical angle compared to level locomotion (α_0_≈43°, α_0_≈38°, α_0_≈39°, and α_0_≈36° for level, 1 cm, 2.5 cm, 5 cm, respectively). Like the trailing leg, the leading leg was significantly less retracted during stance compared to level locomotion. Differences between different steps heights were not significant (Fig. 2 and Table 3).

**Table 3.** Mean, median, max, min values and multiple comparisons for the effective leg during level and step locomotion. For the trailing limb, analyses were performed around TO (TO ± 4%). For the leading limb, at late stance (85% of the stride ± 4%).

|  |  | leg | step up |  |  | step down |  |  | level |
| --- | --- | --- | --- | --- | --- | --- | --- | --- | --- |
|  |  |  | 1 cm | 2.5 cm | 5 cm | 1 cm | 2.5 cm | 5 cm |  |
| leg length [m] | n | tr | 81 | 138 | 144 | 29 | 117 | 80 | 198 |
|  |  | le | 83 | 130 | 118 | 5 | 108 | 54 | 252 |
|  | mean +/- sd | tr | 0.103 +/- 0.005 | 0.108 +/- 0.01 | 0.139 +/- 0.012 | 0.104 +/- 0.013 | 0.107 +/- 0.005 | 0.08 +/- 0.008 | 0.094 +/- 0.005 |
|  |  | le | 0.096 +/- 0.01 | 0.102 +/- 0.006 | 0.108 +/- 0.007 | 0.122 +/- 0.001 | 0.11 +/- 0.007 | 0.097 +/- 0.004 | 0.091 +/- 0.005 |
|  | median | tr | 0.103 | 0.107 | 0.141 | 0.108 | 0.107 | 0.077 | 0.093 |
|  |  | le | 0.1 | 0.104 | 0.109 | 0.122 | 0.111 | 0.095 | 0.091 |
|  | max | tr | 0.111 | 0.126 | 0.155 | 0.123 | 0.117 | 0.096 | 0.107 |
|  |  | le | 0.111 | 0.112 | 0.116 | 0.123 | 0.121 | 0.105 | 0.11 |
|  | min | tr | 0.091 | 0.078 | 0.104 | 0.086 | 0.097 | 0.065 | 0.081 |
|  |  | le | 0.078 | 0.086 | 0.092 | 0.121 | 0.096 | 0.092 | 0.081 |
|  | comp | tr | 1 vs lev (****) | 2.5 vs lev (****)<br>2.5 vs 1 (**) | 5 vs lev (****)<br>5 vs 2.5 (****)<br>5 vs 1 (****) | 1 vs lev (***) | 2.5 vs lev (****)<br>2.5 vs 1 (n.s.) | 5 vs lev (****)<br>5 vs 2.5 (****)<br>5 vs 1 (****) |  |
|  |  | le | 1 vs lev (****) | 2.5 vs lev (****)<br>2.5 vs 1 (***) | 5 vs lev (****)<br>5 vs 2.5 (****)<br>5 vs 1 (****) | 1 vs lev (****) | 2.5 vs lev (****)<br>2.5 vs 1 (**) | 5 vs lev (****)<br>5 vs 2.5 (****)<br>5 vs 1 (****) |  |
| Leg angle ($\alpha$) [°] | n | tr | 81 | 129 | 144 | 36 | 133 | 80 | 198 |
|  |  | le | 83 | 121 | 118 | 18 | 126 | 54 | 252 |
|  | mean +/- sd | tr | 89.1 +/- 10.5 | 96.3 +/- 11.8 | 100.5 +/- 5.7 | 103.6 +/- 19.5 | 82.4 +/- 14.6 | 106.2 +/- 15.7 | 108.2 +/- 10.7 |
|  |  | le | 85.7 +/- 5.8 | 86.1 +/- 8.3 | 84.6 +/- 4.4 | 94 +/- 5.3 | 79.2 +/- 9.4 | 81.4 +/- 5.4 | 88.7 +/- 8.5 |
|  | median | tr | 89.7 | 98.4 | 100.2 | 106.1 | 79.8 | 107 | 110.9 |
|  |  | le | 86.4 | 86.8 | 84.6 | 95.8 | 79.2 | 81.2 | 90.1 |
|  | max | tr | 105.7 | 120.3 | 118.7 | 130.5 | 113.2 | 137.4 | 121.5 |
|  |  | le | 97 | 106.3 | 95.2 | 99.6 | 96.2 | 92.3 | 103.1 |
|  | min | tr | 71.4 | 64.9 | 89.6 | 68 | 52.6 | 62 | 69.7 |
|  |  | le | 73.4 | 71.6 | 75.5 | 82.8 | 64.7 | 71.3 | 59.7 |
|  | comp | tr | 1 vs lev (****) | 2.5 vs lev (****)<br>2.5 vs 1 (**) | 5 vs lev (****)<br>5 vs 2.5 (n.s.)<br>5 vs 1 (****) | 1 vs lev (n.s.) | 2.5 vs lev (****)<br>2.5 vs 1 (****) | 5 vs lev (n.s.)<br>5 vs 2.5 (****)<br>5 vs 1 (n.s.) |  |
|  |  | le | 1 vs lev (**) | 2.5 vs lev (*)<br>2.5 vs 1 (n.s.) | 5 vs lev (****)<br>5 vs 2.5 (n.s.)<br>5 vs 1 (n.s.) | 1 vs lev (*) | 2.5 vs lev (****)<br>2.5 vs 1 (****) | 5 vs lev (****)<br>5 vs 2.5 (n.s.)<br>5 vs 1 (****) |  |

n is the number of points used for multiple comparisons. Significance codes: '\*\*\*\*' ( $p < 0.0001$ ); '\*\*\*' ( $p < 0.001$ ); '\*\*' ( $p < 0.01$ ); '\*' ( $p < 0.05$ ); n.s. (non-significant). tr: trailing limb, le: leading limb. TO: toe-off.

The aperture angle between leading and trailing legs at TD was generally not affected by step height and remained not significantly different from the mean values (φ≈ 53) obtained during level locomotion (p-value > 0.05). Taken together, these observations suggest that effective leg kinematics observed during level locomotion are generally preserved when stepping up onto obstacles.

#### Stepping down, trailing leg

Step related strategies were observed for the trailing leg at the level of the effective leg. Birds negotiating 1 cm drops displayed a compression-extension pattern that diverged from the pattern they exerted during level locomotion and from the monotonic compression displayed when they faced 2.5 cm and 5 cm steps. Stance time was increased with step drop height. Leg compression was significantly larger at TO for 5 cm steps as compared to the other drop conditions.

The trailing leg’s angle of attack (α_0_) was not related to the height of the step-down, and it was similar to the α_0_ observed for level locomotion. For the smallest and largest drops, the trajectory of the effective leg angle was very similar to that observed during level locomotion. For moderate perturbations, the effective leg angle was substantially less retracted during stance. (Fig. 2, Table 3). After TO the leg angle returned to the values observed during level locomotion.

#### Stepping down, leading leg

There were clear adaptations in effective leg kinematics for the leg that stepped on the lowered substrate. The effective leg length at TD for 5 cm step perturbation was significantly shorter than the leg length at TD for 1 cm and 2.5 cm step perturbations (in both cases p-value < 0.0001, see Table 2). During stance, the effective leg was compressed until TO and the effective leg length reached similar values to those observed during level locomotion.

Similarly, effective leg angles were altered during step down locomotion for the leading leg. At TO (elevated substrate) the angle of the effective leg stepping onto the lowered subtract was steeper as compared to level locomotion (2.5 cm: α_TO_≈89°, 5 cm: α_TO_≈87°). Retraction period was prolonged during drops (Table 1). Therefore, the effective leg angle significantly more retracted at TD compared to level locomotion (α_0_≈42°, α_0_≈50°, α_0_≈54°, and α_0_≈53° for level, 1 cm, 2.5 cm, 5 cm, respectively).

The aperture angle between leading and trailing legs was adapted to the drop height. For 1 cm step, the aperture angle increased before TD especially after the level height was crossed. Conversely, for 2.5 cm and 5 cm drops, the aperture angle was on average below the mean value obtained at level locomotion (p-value < 0.0001, respectively p-value < 0.01). Quails adapted the angle between legs after the point at which level height was crossed (Fig. 2).

These observations suggest that effective leg kinematics were substantially altered during step down locomotion.

### Joint angles

The previous section described how effective leg kinematics were altered during uneven locomotion. In this section, we describe how the kinematics of individual, elemental joints were altered. Quail joint angles during level locomotion were previously published ^3^, and therefore, will not be reported here. The influence of the disturbances on the hip angle will be described in the section on 3D hip angles.

#### *Stepping up, trailing limb* (Fig. 3, left column, rows 1 to 3)

To negotiate 1 cm steps, quails used a more flexed INT angle as compared to level locomotion. 2.5 cm perturbations did not induce substantial changes in most joint kinematics. The only exception was the TMP, which is more flexed at TD. To negotiate 5 cm steps the knee and the INT joints were significantly more extended, and the TMP was more flexed during stance. After TO, the knee was kept more extended during the early swing phase. Note that the bouncing behavior observed in the INT almost vanishes when facing 5 cm step up perturbations.

**Figure 3.**
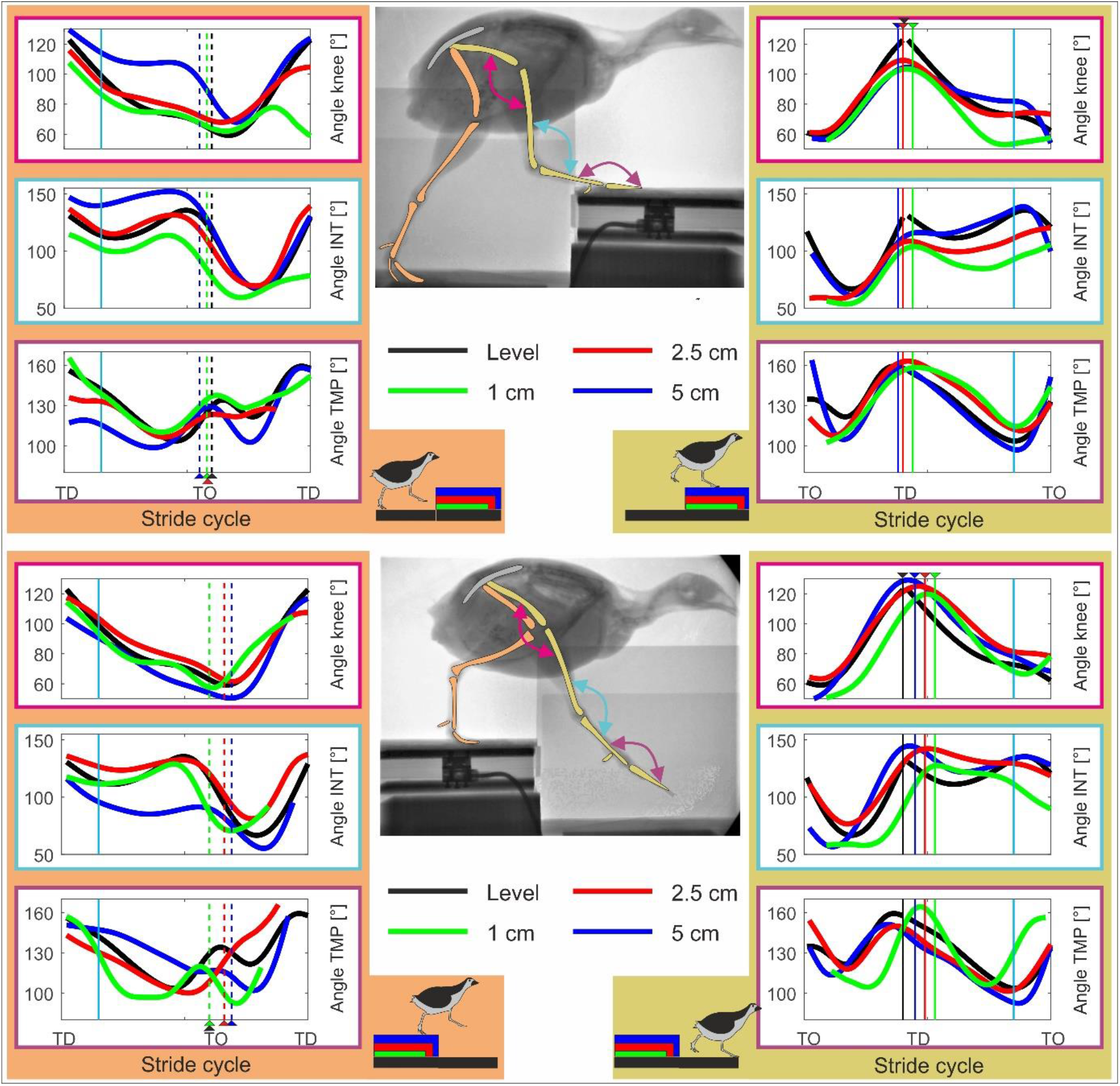
Joint angles. Knee, intertarsal (INT) and tarsometatarsal-phalangeal (TMP) joint angles during level (black) and step locomotion (1cm: green, 2.5 cm: red, 5 cm: blue) in the quail. Rows 1 to 3 level vs. step up locomotion. Rows 4 to 6 level vs step down locomotion. Left: trailing limb (stride i-1), right: leading limb (stride i). Curves display mean values. Black, blue, red, green dashed lines indicate toe-off (TO), while solid lines touch down (TD). Cyan solid lines indicate 15% and 85% of the stride.

#### *Stepping up, leading limb* (Fig 3, right column, rows 1 to 3)

In the elevated substrate, the quails displayed a more flexed knee and INT at TD for all perturbations. During stance on the step, the joint patterns for 1 cm and 2.5 cm steps quails displayed a more flexed INT, together with a more extended TMP compared to the patterns observed for 5cm steps.

#### *Stepping down, trailing limb* (Fig. 3, left column, rows 4 to 6)

When negotiating 1 cm steps, the flexion-extension pattern for the TMP changed. Note that during stance there was a larger flexion up to midstance, followed by an extension in the late stance. After TO, a second more marked flexion extension was exhibited. For 2.5 cm drops, quails displayed a stiffer INT, perhaps to vault downwards. More marked differences in all joints were observed for 5 cm steps. Under this test condition, knee and INT joints exhibited significantly larger flexion at TD and during stance. After TO, knee and INT and were kept more flexed.

#### *Stepping down, leading limb* (Fig. 3, right column, rows 4 to 6)

The leg that stepped in the lowered substrate, displayed step related adaptations before and after TD. Before TD, changes were observed mainly in the distal joints. 1 cm drops increased joint flexion in the first half of the swing phase but did not induce significant changes at TD related to level locomotion. 2.5 cm and 5 cm drops did not substantially influence joint swing patterns but affected joint angles at TD (significantly more extended for the knee and INT and significantly more flexed for the TMP, see Table 4). After TD, the INT was further flexed for 1 cm and 2.5 cm drops until TO. The INT for 5 cm and the TMP for 1 cm drops displayed a rebound behavior (flexion-extension pattern). For 2.5 cm and 5 cm drops, TMP patterns were like those observed for level locomotion, but the joints were kept more flexed until late stance. Adaptations in limb kinematics display a shift from a dynamic to a safety guided gait program as perturbation drop height increases.

**Table 4.**
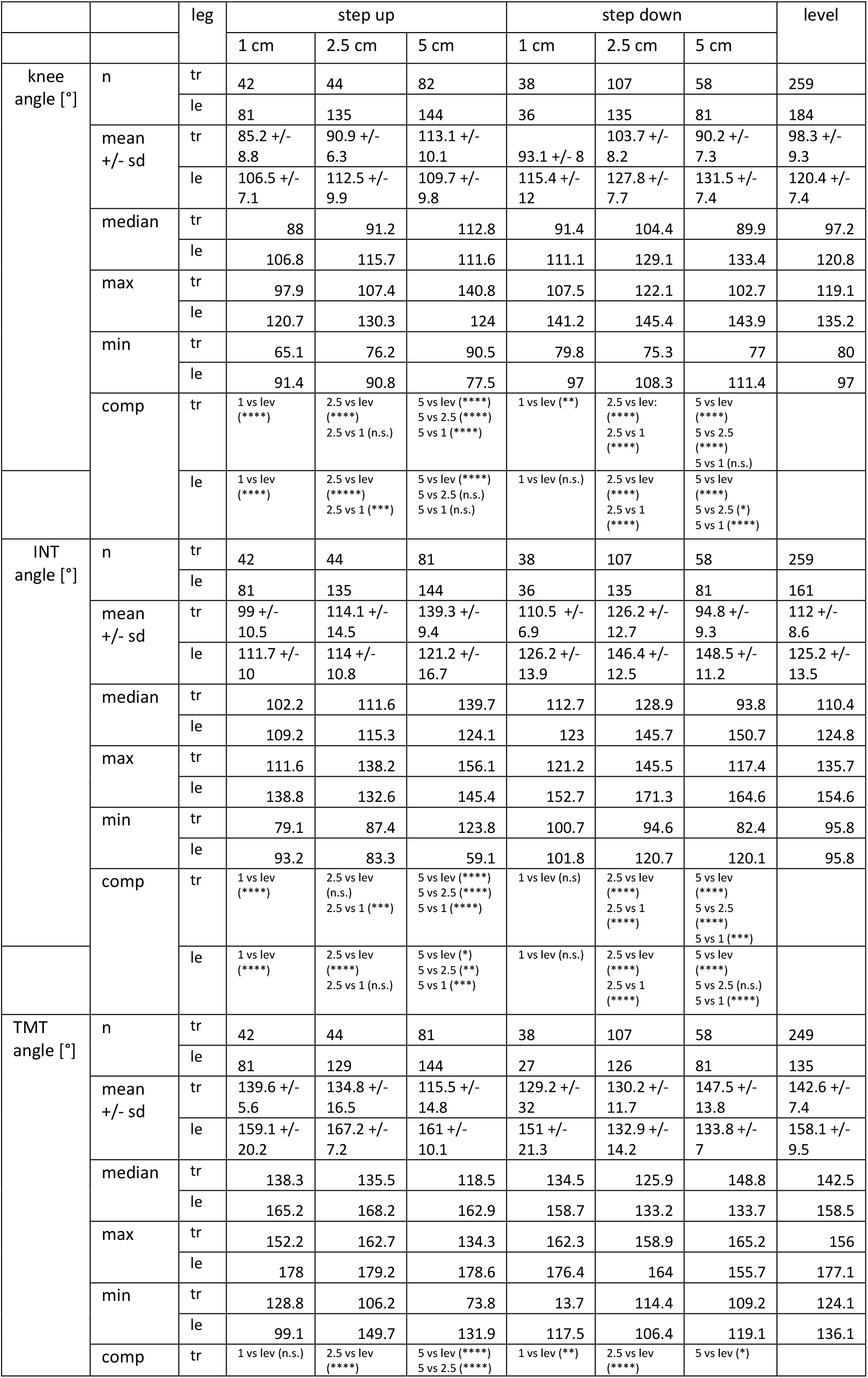
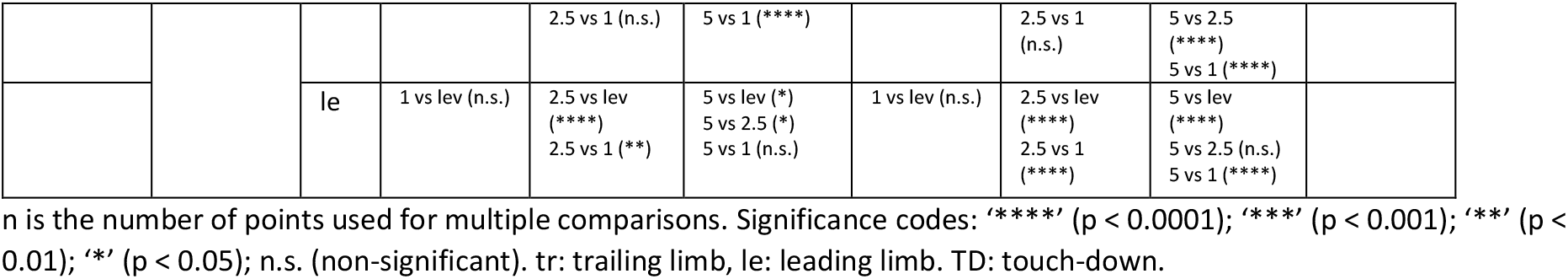
Mean, median, max, min values and multiple comparisons between joint angles during level and step locomotion. For the trailing limb, analyses were performed at early stance (15% of the stance ± 4%). For the leading limb, around TD (TD ± 4%).

**Table 5.**
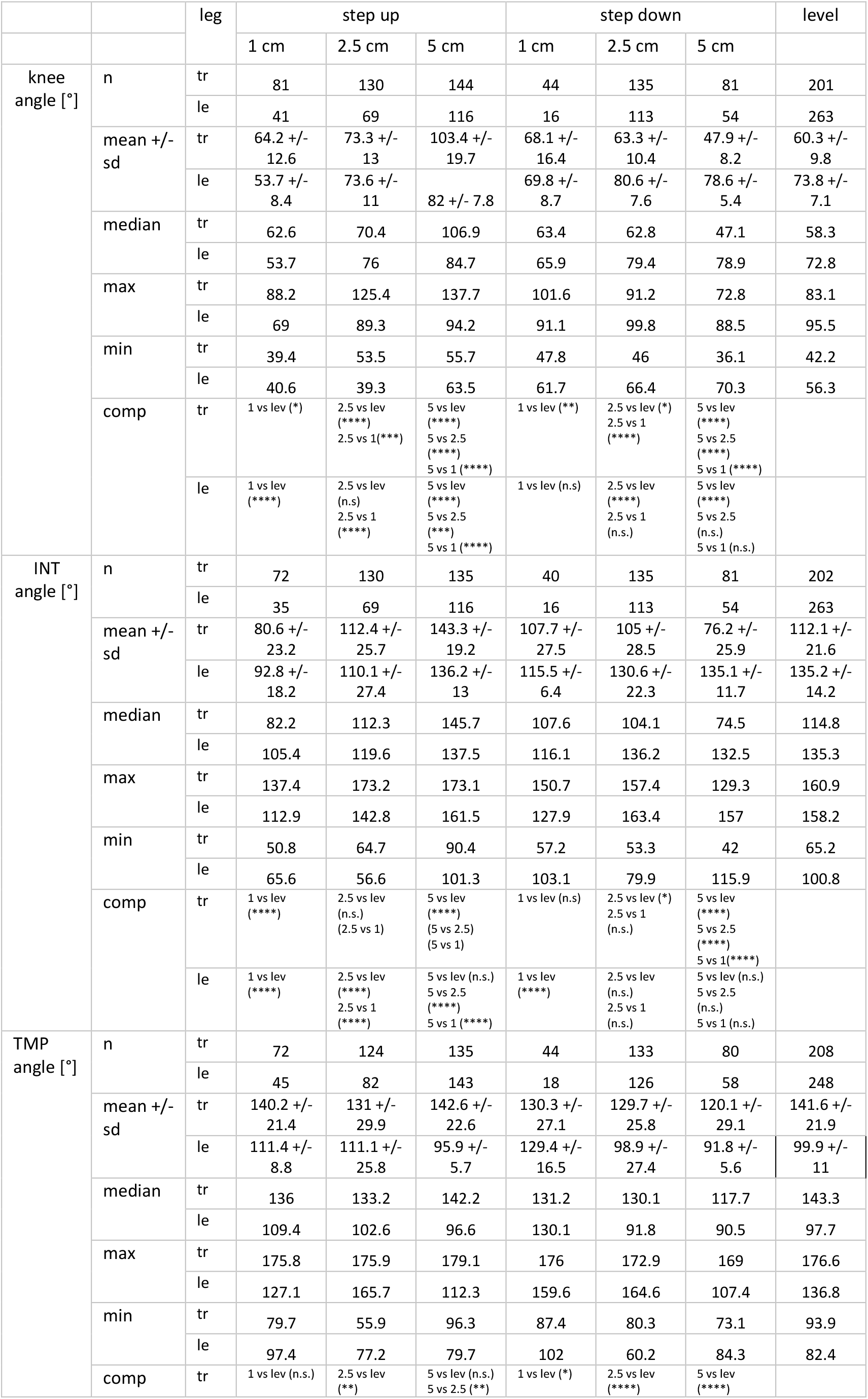
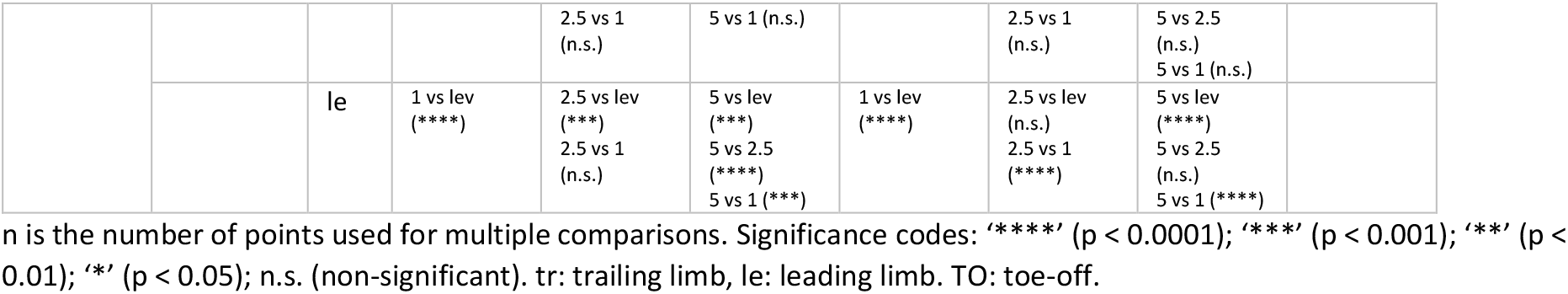
Mean, median, max, min values and multiple comparisons between joint angles during level and step locomotion. For the leading limb, analyses were performed at late stance (85% of the stance ± 4%). For the trailing limb, around TO (TO ± 4%).

### 3D-kinematics of the whole leg

This section describes the three-dimensional kinematics of the whole leg relative to the pelvis during level and step locomotion (see Fig. 4). Under the assumption that both knee and intertarsal joints work as revolute joints the whole leg approximates three-dimensional hip kinematics. Note that because the z-axis was aligned with the segment from hip to knee, rotation about y-axis (β) reflects flexion/extension between femur and pelvis, rotations about z-axis (γ) reflect hip ab-adduction, while rotations about the x-axis (α) reflect femoral axial rotations, resulting in the lateromedial rotation of the whole leg. α = β = γ = 0° indicates that the whole leg and the pelvis coordinate systems are aligned. However, in this zero-pose, the pelvis and femur are orthogonal to each in the sagittal plane. Therefore, we used β + 90° to represent hip flexion/extension in Fig. 4 and Tables 6 and 7. In the following, level locomotion is first described in detail. Step locomotion is discussed when there is a difference from level locomotion.

**Figure 4.**
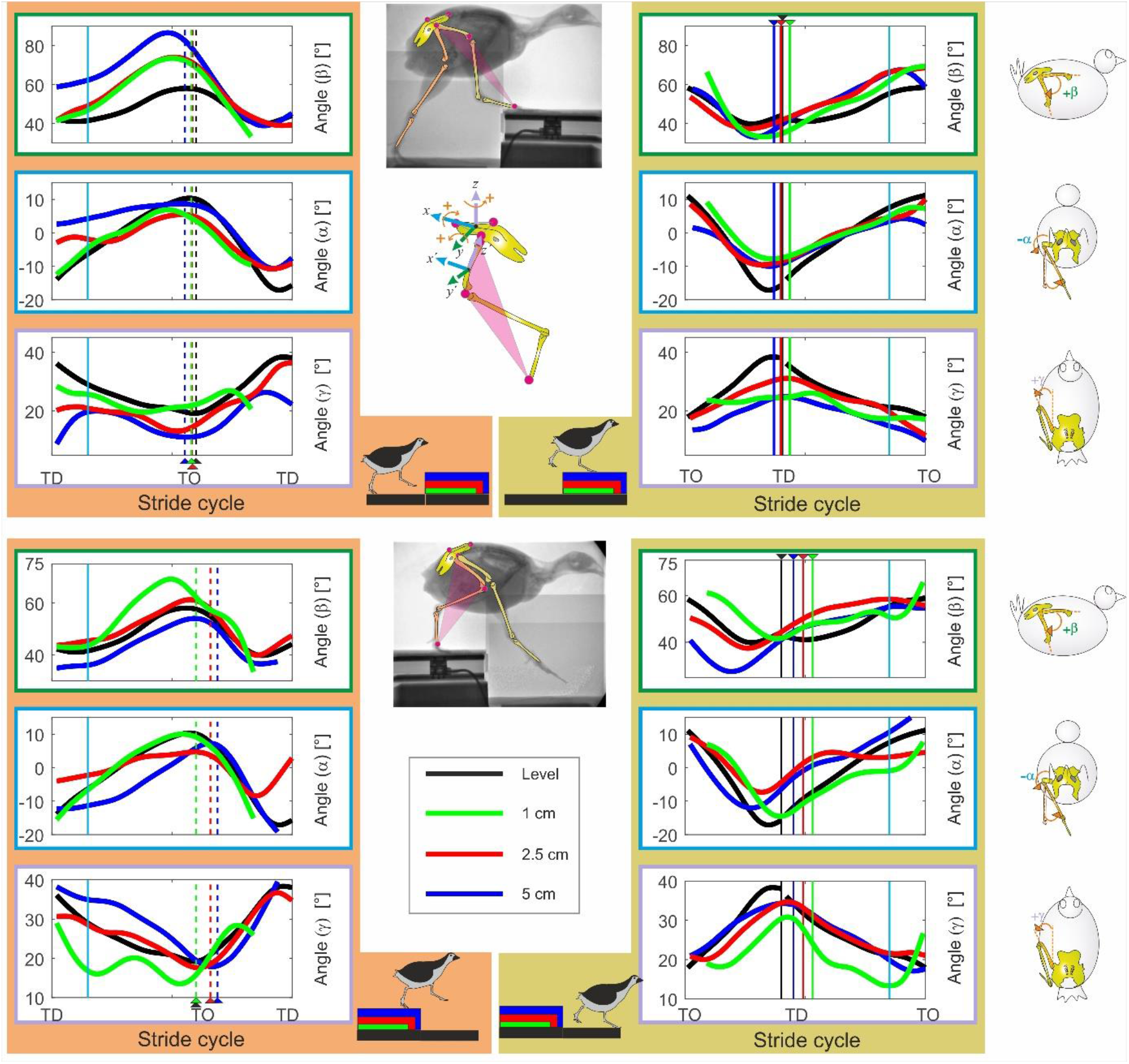
Whole leg three-dimensional rotations in the quail. Motions were measured relative to the pelvis. Level (black) and step locomotion (1cm: green, 2.5 cm: red, 5 cm: blue). Accepting that the knee, the intertarsal and the tarsometatarsal-phalangeal joints work mainly as revolute joints, the plane describing the whole-leg displays the three-dimensional hip control. Curves display mean values. Rows 1 to 3 display level vs. step-up locomotion. Rows 4 to 6 level vs step-down locomotion. Left: trailing limb (steps before the vertical perturbation, stride i-1), right: leading leg (steps after the vertical perturbation, stride i). Rows 1 & 4: hip flexion extension, negative values indicate flexion. Rows 2 and 5: lateromedial rotation. Positive values indicate that the distal point of the whole leg moves laterally with respect to the hip. Rows 3 and 6: Ad-abduction of the whole leg. Black, blue, red, green dashed lines indicate toe-off (TO), while solid lines touch down (TD). Cyan solid lines indicate 15% and 85% of the stride.

**Table 6.**
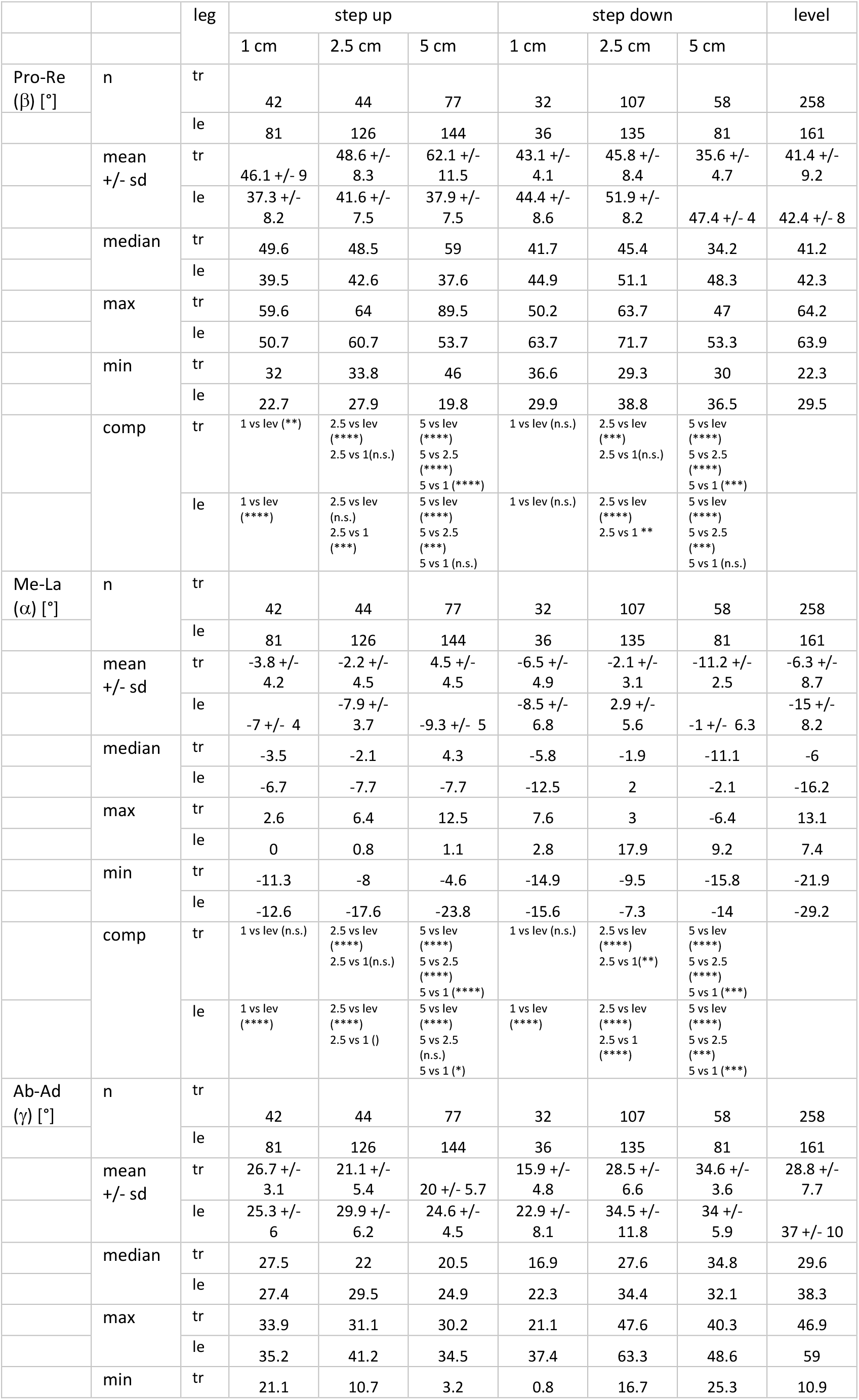
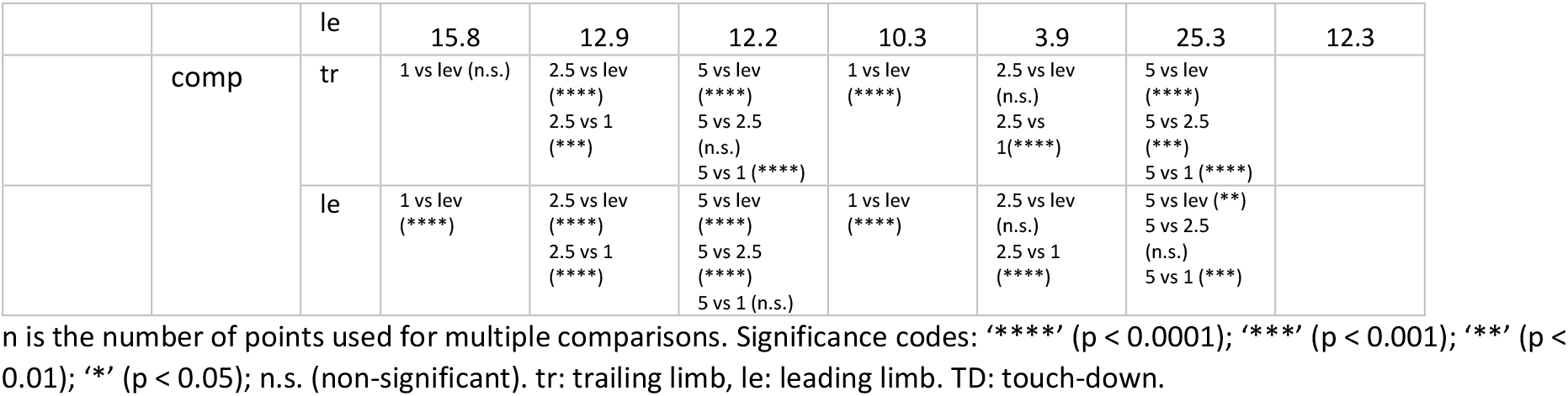
Mean, median, max, min values and multiple comparisons between hip cardan angles during level and step locomotion. For the trailing limb, analyses were performed at early stance (15% of the stance ± 4%). For the leading limb, around TD (TD ± 4%).

**Table 7.**
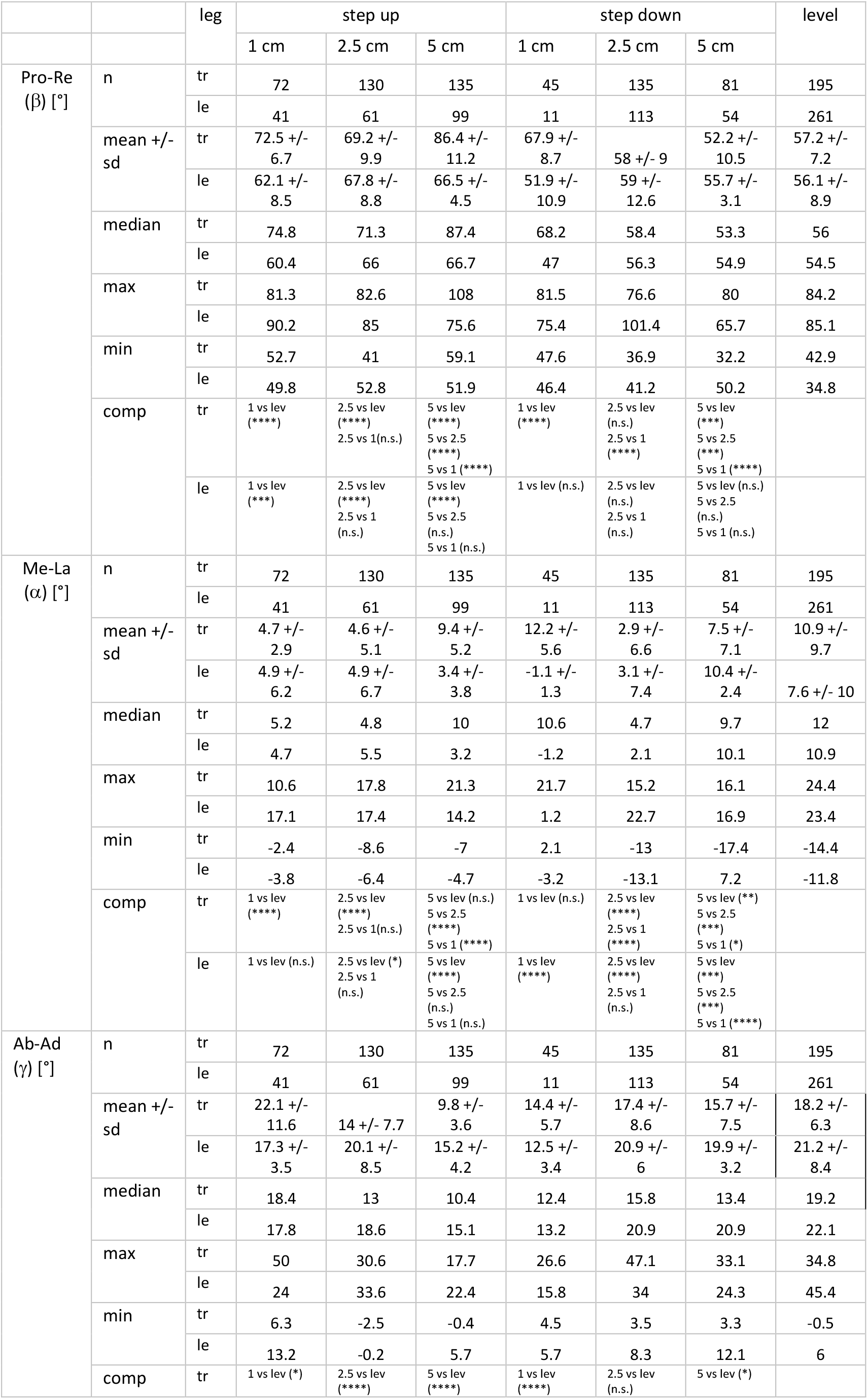
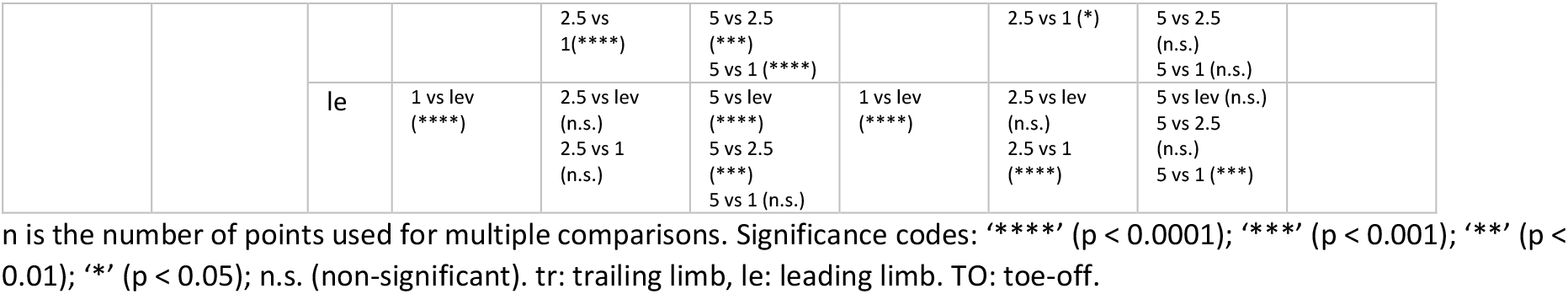
Mean, median, max, min values and multiple comparisons between hip cardan angles during level and step locomotion. For the leading limb, analyses were performed at late stance (85% of the stance ± 4%). For the trailing limb, around TO (TO ± 4%).

**Table 8.**
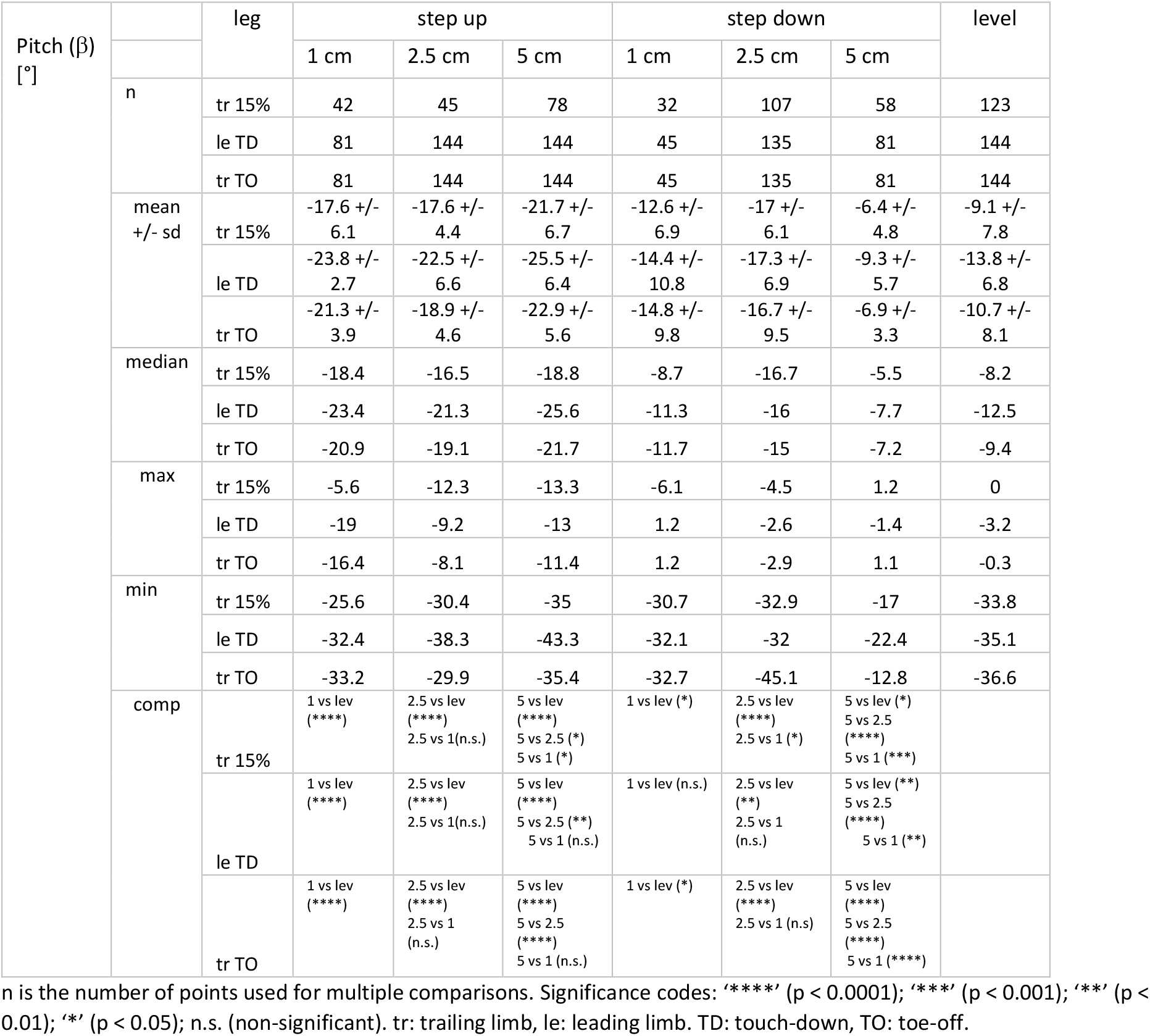
Mean, median, max, min values and multiple comparisons between pelvic pitch angles during level and step locomotion. For the trailing limb, analyses were performed at early stance (15% of the stance ± 4%) and around TO (TO ± 4%). For the leading limb, around TD (TD ± 4%).

**Table 9.**
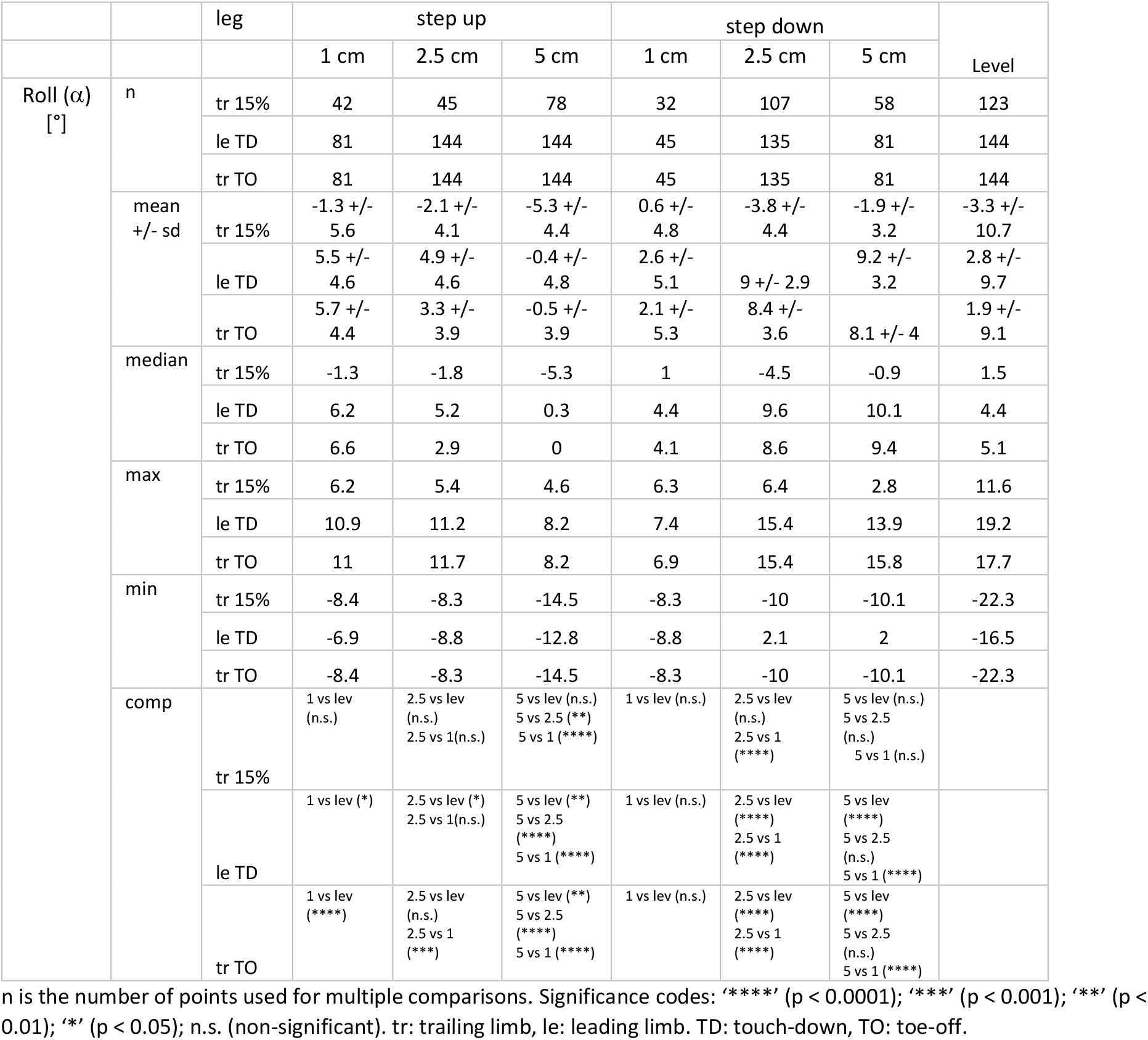
Mean, median, max, min values and multiple comparisons between pelvic roll angles during level and step locomotion. For the trailing limb, analyses were performed at early stance (15% of the stance ± 4%) and around TO (TO ± 4%). For the leading limb, around TD (TD ± 4%).

**Table 10.**
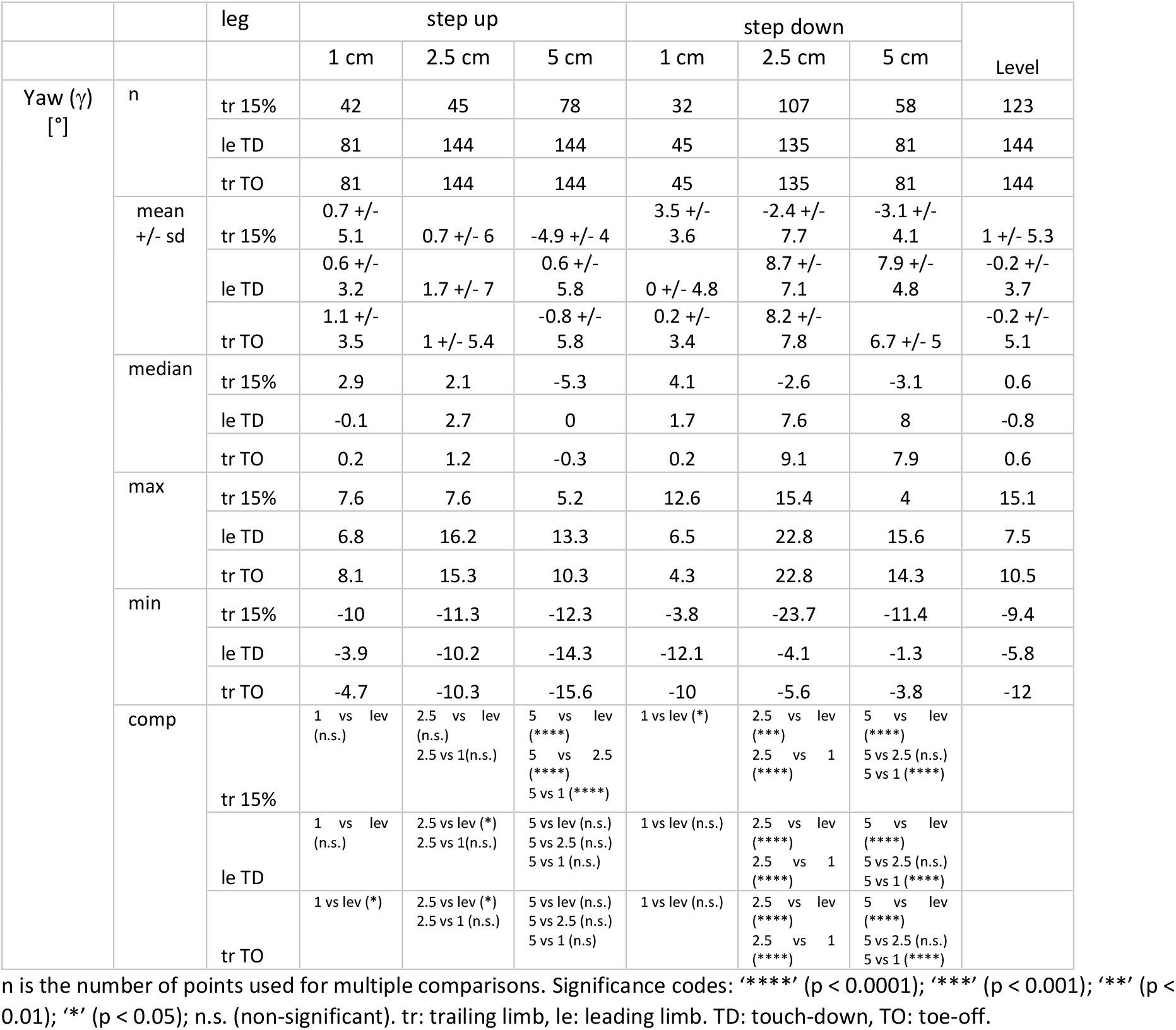
Mean, median, max, min values and multiple comparisons between pelvic yaw angles during level and step locomotion. For the trailing limb, analyses were performed at early stance (15% of the stance ± 4%) and around TO (TO ± 4%). For the leading limb, around TD (TD ± 4%).

|  |  | leg | step up |  |  | step down |  |  | Level |
| --- | --- | --- | --- | --- | --- | --- | --- | --- | --- |
|  |  |  | 1 cm | 2.5 cm | 5 cm | 1 cm | 2.5 cm | 5 cm |  |
| Yaw ($\gamma$)<br>[°] | n | tr 15% | 42 | 45 | 78 | 32 | 107 | 58 | 123 |
|  |  | le TD | 81 | 144 | 144 | 45 | 135 | 81 | 144 |
|  |  | tr TO | 81 | 144 | 144 | 45 | 135 | 81 | 144 |
|  | mean<br>+/- sd | tr 15% | 0.7 +/-<br>5.1 | 0.7 +/- 6 | -4.9 +/- 4 | 3.5 +/-<br>3.6 | -2.4 +/-<br>7.7 | -3.1 +/-<br>4.1 | 1 +/- 5.3 |
|  |  | le TD | 0.6 +/-<br>3.2 | 1.7 +/- 7 | 0.6 +/-<br>5.8 | 0 +/- 4.8 | 8.7 +/-<br>7.1 | 7.9 +/-<br>4.8 | -0.2 +/-<br>3.7 |
|  |  | tr TO | 1.1 +/-<br>3.5 | 1 +/- 5.4 | -0.8 +/-<br>5.8 | 0.2 +/-<br>3.4 | 8.2 +/-<br>7.8 | 6.7 +/- 5 | -0.2 +/-<br>5.1 |
|  | median | tr 15% | 2.9 | 2.1 | -5.3 | 4.1 | -2.6 | -3.1 | 0.6 |
|  |  | le TD | -0.1 | 2.7 | 0 | 1.7 | 7.6 | 8 | -0.8 |
|  |  | tr TO | 0.2 | 1.2 | -0.3 | 0.2 | 9.1 | 7.9 | 0.6 |
|  | max | tr 15% | 7.6 | 7.6 | 5.2 | 12.6 | 15.4 | 4 | 15.1 |
|  |  | le TD | 6.8 | 16.2 | 13.3 | 6.5 | 22.8 | 15.6 | 7.5 |
|  |  | tr TO | 8.1 | 15.3 | 10.3 | 4.3 | 22.8 | 14.3 | 10.5 |
|  | min | tr 15% | -10 | -11.3 | -12.3 | -3.8 | -23.7 | -11.4 | -9.4 |
|  |  | le TD | -3.9 | -10.2 | -14.3 | -12.1 | -4.1 | -1.3 | -5.8 |
|  |  | tr TO | -4.7 | -10.3 | -15.6 | -10 | -5.6 | -3.8 | -12 |
|  | comp | tr 15% | 1 vs lev<br>(n.s.) | 2.5 vs lev<br>(n.s.)<br>2.5 vs 1(n.s.) | 5 vs lev<br>(****)<br>5 vs 2.5<br>(****)<br>5 vs 1 (****) | 1 vs lev (*) | 2.5 vs lev<br>(***)<br>2.5 vs 1<br>(****) | 5 vs lev<br>(****)<br>5 vs 2.5 (n.s.)<br>5 vs 1 (****) |  |
|  |  | le TD | 1 vs lev<br>(n.s.) | 2.5 vs lev (*)<br>2.5 vs 1(n.s.) | 5 vs lev (n.s.)<br>5 vs 2.5 (n.s.)<br>5 vs 1 (n.s.) | 1 vs lev (n.s.) | 2.5 vs lev<br>(****)<br>2.5 vs 1<br>(****) | 5 vs lev<br>(****)<br>5 vs 2.5 (n.s.)<br>5 vs 1 (****) |  |
|  |  | tr TO | 1 vs lev (*) | 2.5 vs lev (*)<br>2.5 vs 1 (n.s.) | 5 vs lev (n.s.)<br>5 vs 2.5 (n.s.)<br>5 vs 1 (n.s.) | 1 vs lev (n.s.) | 2.5 vs lev<br>(****)<br>2.5 vs 1<br>(****) | 5 vs lev<br>(****)<br>5 vs 2.5 (n.s.)<br>5 vs 1 (****) |  |

n is the number of points used for multiple comparisons. Significance codes: '\*\*\*\*' (p < 0.0001); '\*\*\*' (p < 0.001); '\*\*' (p < 0.01); '\*' (p < 0.05); n.s. (non-significant). tr: trailing limb, le: leading limb. TD: touch-down, TO: toe-off.

#### *Level locomotion, hip flexion-extension* (β)

At TD, the hip joint is flexed about 42°. After a small flexion due to weight transfer, the hip joint extends 17° until TO. After TO the leg protracts, flexing the hip joint up to 85% of swing. In the late swing phase, the whole leg retracts until TD.

#### *Level Locomotion, lateromedial control of the whole leg* (α)

At TD the whole leg was medially oriented (α ≈ −14°). During stance, the leg was rotated laterally until TO to an angle of approx. α = 11°. During swing the distal point of the whole leg was rapidly rotated medially.

#### *Level Locomotion, whole leg (femoral) ab-adduction* (γ)

hip ab-adduction curves show a half-sine pattern. At TD the whole leg was abducted about 36°. Abduction was reduced during stance to 18° at TO. After TO the leg was abducted up to TD.

#### *Stepping up, trailing limb* (Fig. 4, left column, rows 1 to 3)

Step height had a significant influence on hip flexion-extension. At TD, quails facing 5 cm steps exhibited significant larger hip extension. As stance phase progressed, the hip joint was significantly more extended during stepping up than during level locomotion (p-values= 0.0042, 0.00003, and 0 for 1cm, 2.5 cm, and 5 cm, respectively). However, 1 cm and 2.5 cm steps induced, on average, similar hip extension patterns (p-value > 0.05) but significantly different from 5 cm (i.e., quails displayed a two-step strategy to negotiate vertical perturbation). Mediolateral hip control was also influenced by step height. At TD, 2.5 cm and 5 cm step ups induced a more vertical orientation of the whole leg, and at TO the whole leg was less laterally oriented than during level locomotion. During step-up locomotion the whole leg was less abducted. While quails facing 5 cm steps decreased abduction in similar way as when they negotiated 2.5 cm steps, for coping with 1 cm steps they kept adduction similar to the abduction observed during level locomotion. After TO, quails facing 2.5 cm and 5 cm steps increased abduction, approaching values observed during level locomotion. However, for 5 cm steps, quails maintained a persistent hip adduction in the late swing.

#### *Stepping up, leading limb* (Fig. 4, right column, rows 1 to 3)

Flexion-extension patterns in the elevated step are similar in shape to those observed for level locomotion. However, the quail stepped with a more flexed hip after negotiating 1 cm and 5 cm steps. After TD, the quail exhibited comparative larger hip extensions compared to level locomotion (see Table 7 for comparison at late stance). In contrast, the quail reduced both mediolateral rotations and ab-adduction during the swing phase before stepping on the elevated substrate. At TD on the elevated substrate, the leading whole leg was significantly less abducted and more vertically oriented compared to level locomotion. After the early stance phase, mediolateral motion differences between step and level locomotion lessened. For 1 cm steps, the abduction of the whole leg stayed around γ = 20°.

#### *Stepping down, trailing limb* (Fig. 4, left column, rows 4 to 6)

Quails facing 1 cm visible drops displayed larger hip extension after midstance. This can be explained by the tendency of the subjects to switch to aerial running when negotiating this type of perturbation. 2.5 cm drops did not induce major changes in the flexion-extension patterns of the hip. When negotiating 5cm drops, the hip joint was significantly more flexed than during level locomotion.

The response of the mediolateral hip control for 1 cm and 2.5 cm was similar to those observed during step-up perturbations. For 5 cm drops, the leg was medially oriented at TD like observed during level locomotion and straightening of the leg during stance was more gradual.

The abduction of the leg increased with drop height. When quails faced 1 cm steps, adduction of the whole leg was reduced with respect to level locomotion. When they negotiated 2.5 cm steps, abduction was on average similar to the patterns exhibited during level locomotion (see Table 6), while for 5 cm drops, the whole leg was kept more abducted during stance.

#### *Stepping down, leading limb* (Fig. 4, right column, rows 4 to 6)

quails started the swing phase using a more extended hip to approach 1 cm drops, and more flexed for 2.5 cm and 5 cm drops. At TD in the lowered substrate, the hip was more extended for 1 cm (not significant), 2.5 cm and 5 cm.

Whole leg medial rotations (femoral outer rotations) were constrained when negotiating 2.5 cm and 5 cm drops. This permitted the quail to step in the lowered substrate with an almost vertically oriented whole leg.

Hip adduction was also reduced during the swing phase. After 1 cm drop, the quail kept their hip more adducted during stance, but close before TO, the hip joint was abducted. After 2.5 cm and 5 cm drops hip adduction behaved like the patterns observed for level locomotion.

### Pelvis

The pelvis/trunk of a quail was controlled as a single body (Fig. 5). Pelvic pitch oscillation frequency was twice the step frequency, across all locomotion conditions Compared to level locomotion, pelvic retroversion increased when the quail negotiated step up perturbations: the pelvis was retroverted about 10° during level and up to 28° during step up locomotion. For visible drops the picture was less clear and was inconsistent across different size drops. Relative to the values obtained for level locomotion, when quails faced 1 cm drops, they increased and then decreased pelvic retroversion after the TD in the lowered substrate. When they faced 2.5 cm drops, they increased pelvic retroversion (mean values oscillated about 20°), and when quails negotiated 5cm drops, they decreased pelvic retroversion.

**Figure 5.**
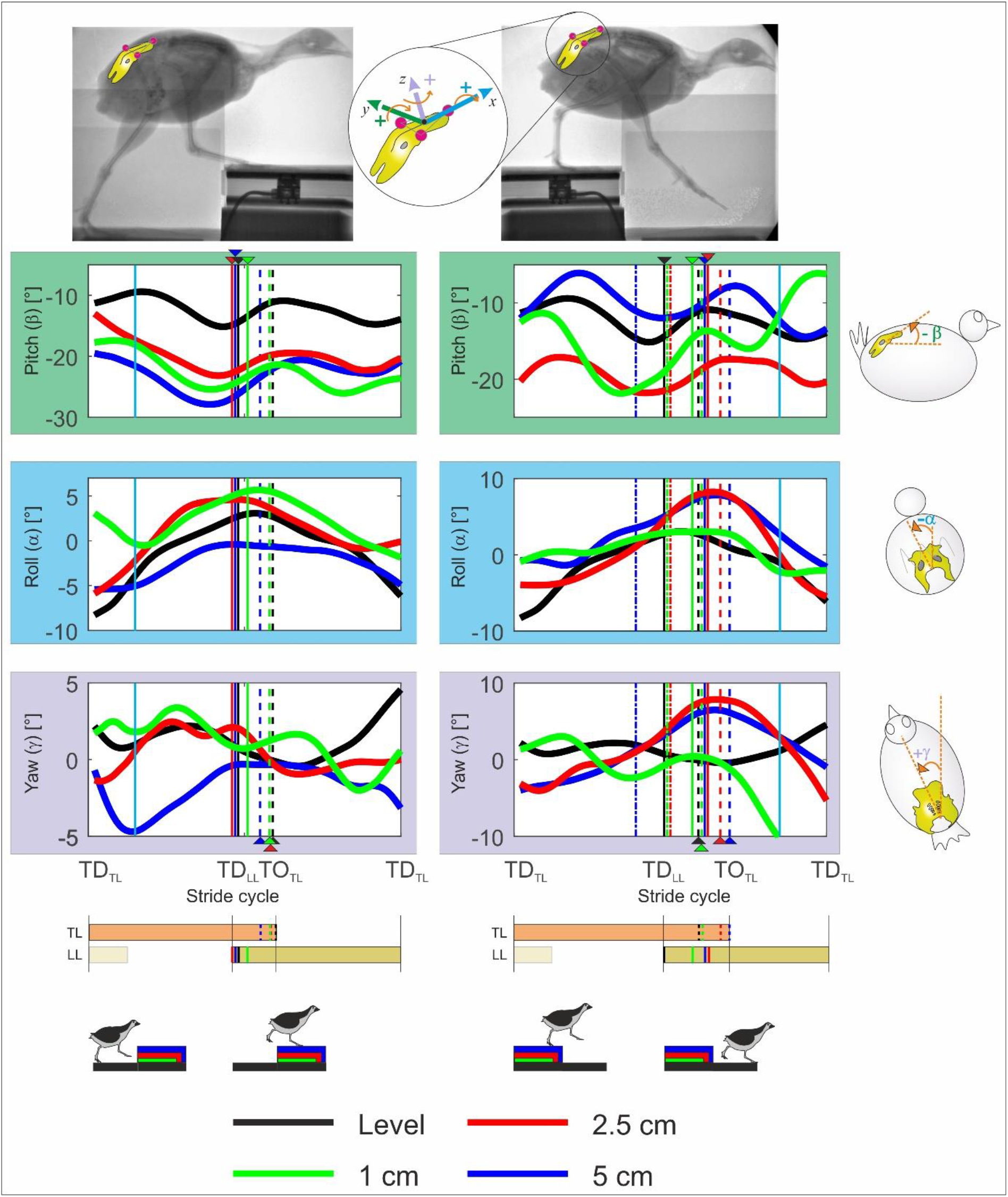
Pelvic three-dimensional rotations during level (black) and step locomotion in the quail. Curves display mean values. Left: step-up locomotion, right: step-down locomotion. For better understanding, we transformed the data to ensure that the trailing limb is always the left leg and the leading leg the right one (see methods). Upper: pelvic pitch, negative values indicate retroversion (trunk is more vertical oriented). Middle: pelvic roll, positive values indicate that the trunk tilts towards the right. Bottom: Pelvic yaw, positive values indicate that the body is directed towards the left. Black, blue, red, green dashed lines indicate toe-off of the contralateral leg (TO), while solid lines touch down (TD). Dot dashed lines indicate when the leg cross level line in step-down perturbations. Cyan solid lines indicate 15% and 85% of the stride. TL: trailing limb, LL: leading limb.

Lateral tilt (roll) was cyclic and counteracted by the leg in contact with the substrate. Pelvic yaw amplitudes were small, but there was a rotation of the pelvis towards the direction of the leg in contact with the ground. To facilitate negotiating larger visible drops, the pelvis (and the trunk) were rotated towards the trailing limb (yaw) and tilted (roll) towards the leading leg. After TD in the lowered substrate, the pelvis (trunk) was reoriented in motion’s direction.

## Discussion

To understand control strategies implemented by any system, it is necessary to characterize how the system responds to external perturbations. In the present work we analyzed the kinematic strategies employed by the common quail to negotiate visible step-up and step-down perturbations of about 10%, 25%, and 50% of the average value of their effective leg length during stance. Our main goal was to uncover leg kinematic changes at different levels of abstraction and how they relate to each other. The highest level of abstraction in our work is found in the effective leg (Fig. 1E). The kinematic analysis of the effective leg characterizes global control goals such as leg length, angle of attack at TD, aperture angle and retraction speed. Note that the effective leg will have two main functions if the dynamics are taken into consideration: a) the axial leg function, which is a time-dependent force function (e.g., spring-damper) and b) the tangential or rotational leg function, which is a time-dependent torque that controls the leg and balances the trunk (e.g., virtual pivot point (VPP) control ^18,35^). Two- and three-dimensional joint kinematics (Figs. 1F and 1D) are representations with less level of abstraction. Because different combinations of joint kinematics can lead to the same effective leg lengths, we expected that their combined analysis would help to infer quail motor control goals on uneven terrains. Thus, we compared the a) effective leg kinematic, b) joint kinematics and c) whole leg (represents hip 3D kinematics, see Fig. 4) and pelvic kinematics for the quail negotiating step-up and step-down perturbations with our previously collected data on quail level ground running ^18^, which is freely available on https://datadryad.org/stash/dataset/doi:10.5061/dryad.jh5h4.

Our results display a complex picture of kinematic strategies before and after TD. In the next sections, we analyze that complex picture by linking our results with the existing knowledge about the interactions between kinematics, dynamics, and muscle activation during level/uneven locomotion. This combined analysis is used to unravel anticipatory and reactive strategies for the negotiation of step perturbations, and to discuss whether those strategies may be governed by simple control goals.

### Stepping up

#### Trailing limb (stride i-1)

In the step before the perturbation (i-1), the trailing effective leg was significantly longer at TD for stepping up than observed during level grounded running. Moreover, the effective leg length significantly increased with step height. The angle of attack at TD was steeper as step height increased. The differences in effective leg length between level locomotion and step locomotion at TD might be explained by the fact that data for level and step locomotion belonged to different quail cohorts. Animals had similar age, but the quail facing steps were heavier. However, longer effective leg length at TD and steeper angle of attack at TD might also indicate a “pre-programmed” control strategy at the global level to negotiate upward steps perhaps producing a shift in the operating locomotion program towards “mixed gaits” ^21^, a periodic change between walking and grounded running steps that might permit birds to adjust their leg to vault towards the elevated substrate ^10^. A more extended leg at TD also would agree with observations in running humans, which adapt their center of mass (CoM) height about 50% of step height in anticipation of stepping onto a visible step ^36,37^. Note that because of neuromuscular delays, vertebrates preset muscle force before TD using posture dependent control ^3,10–12,38^. During stance, the quail also fine-tuned leg length, and leg retraction of the trailing effective leg according to step height (see Fig. 2). This adjustment indicates that visual perception of the upcoming obstacle induced anticipatory changes in leg loading during stance. One can hypothesize that the goal of this sensory driven adaptation was to adjust the trajectory of the CoM to reduce the necessity of compensation in the following step.

How was the effective trailing leg length adjusted at the joint level in the step before the perturbation? Our results suggest that the quail used two distinct strategies, depending on the height of the step. For step heights up to 25% of effective leg length, the extension of the hip joint lengthened the leg, while knee and intertarsal joints displayed similar patterns to those observed during level locomotion. For the 5 cm step height (about 50% of effective leg length) both knee and intertarsal joints were extended, while the hip joint extended even more.

Note that during quail level locomotion, the spring-like leg behavior is mostly produced in the INT, while the active flexion of the knee joint controls leg retraction ^3^. However, to negotiate 5 cm steps, the extension of both knee and INT turned the crouched quail leg into a more vertical one. In this leg configuration, the retraction of the leg is produced by hip extension. Thus, to vault the CoM onto the obstacle, the avian leg was controlled in a similar manner as humans and animals, which have a more stiff and extended leg design.

Thus, the “zig-zag” configuration of the femur, the tibiotarsus, and the tarsometatarsus is abandoned to negotiate larger vertical perturbations (see the trailing the limb configuration superimposed to the X-ray picture in Fig. 3). The enclosed joints are spanned by mono- and bi-articular muscles with the latter enforcing a parallel mechanism, the so called pantograph leg ^39,40^. Gordon and colleagues ^9^ reported significant larger activations for muscles *M. flexor cruris lateralis pelvica* (FCLP, hip extensor, knee flexor, possible hip abductor), *M. gastrocnemius pars lateralis* (GL, ankle extensor, knee flexor), *M. gastrocnemius pars medialis* (GM, ankle extensor, knee flexor/extensor), *M. flexor perforatus digiti* III (FPPD3, ankle extensor, digital flexor), and *M. femorotibialis lateralis* (FTL, mono-articular knee extensor) in the step prior to a step-up perturbation. These activation profiles are consistent with the control of the extension in the hip joint, the knee and the INT in the quail. In addition, the larger activation of FCLP correlates also with the reduced hip adduction in the quail when negotiating 5 cm step-up perturbations. At the neuronal level this shift in leg behavior might be induced by changed muscle synergies via higher locomotor center signals based on visual perception.

#### Leading leg towards and on the elevated substrate (stride i)

When the leading limb was swung towards the elevated substrate, the quail controlled the aperture angle between legs as described for level locomotion ^4^. In the late swing the aperture angle was kept constant at a φ≈53° despite step height. Thus, the late swing retraction and the angle of attack of the leading leg were mainly controlled by the retraction of the trailing leg as hypothesized.

When the leading leg stepped on the elevated substrate, the effective leg length and the angle of attack were similar to those observed in level locomotion. After TD, the effective leading leg kinematics did not markedly differ from those observed during level locomotion. Adaptations of the trailing limb thus permitted the leading limb to touch down on the step in similar manner as during level locomotion. This strategy might help to rapidly dissipate the perturbations produced by the vertical step. Empirical evidence has shown that running animals recover steady state behavior two to three steps after an unexpected perturbation ^11,20,41^. Our results suggest that the quail recovered even faster from a visible perturbation (mostly one step), as described previously for other birds ^9–11^.

Despite the significant extension of the trailing leg, the leading leg touched down with joints more flexed than during level locomotion. After TD, the hip was rapidly more extended than during level locomotion, and the behavior of the INT shifted from a spring-like mode to an energy supplier (joint extended beyond its angle at TD) as step height increased. Note that at TD, the knee was not used to extend the leg, possibly because larger extensor torques about this joint would increase the horizontal GRF, breaking the retraction of the leg. Even so, the flexion of the knee was controlled during stance when negotiating the largest step heights, so that the knee-joint angle returned slowly to the value exhibited during unrestricted locomotion. The increased extensor activity of the FTL muscle observed after the Guinea Fowl stepped on an elevated substrate, might be consistent with our observations ^9^.

In summary, even when the trailing leg extension might have reduced the necessity of reactive control, changes in loading of the leading leg might be necessary to compensate for the more flexed joints at TD.

### Stepping down

#### Trailing limb (stride i-1)

When the quail negotiated drops of about 10% of effective leg length, they used aerial phases to rapidly overcome the perturbation. To introduce aerial phases, the operation of the trailing leg was shifted towards spring-like behavior (more marked rebound, see Fig. 2). At the effective leg level, this change can be produced by reducing effective leg damping and/or inducing an axial extension of the effective leg in the late stance. In both cases, the pronograde virtual pivot point model [PVPP, ^18^] predicts that the axial energy of the system increases. This makes aerial phases more likely to occur. But how are those changes produced at the joint level? As observed for step-up perturbations, hip extension seems to control effective leg extension if legs are kept crouched (c.p. Fig. 2 and Fig. 4). Knee and INT joint kinematics did not display sudden changes compared to level locomotion (Fig. 2). This seems to indicate, following ^3^, that neither retraction angle, nor effective leg stiffness were adapted to negotiate the lowest drop height. Indeed, the trajectories for the retraction angle did not deviate from those observed during level locomotion (see Fig. 2). Note that we did not estimate leg stiffness for this study. To estimate it, it is necessary to combine ground reaction force data together with the effective leg length change ^18^. Thus, our predictions in this respect are educated guesses based on our previous works on dynamics of bipedal level and perturbed locomotion.

When compared with the patterns obtained during level locomotion, the TMP joint displayed a change to a more spring-like function (see Fig. 2). Because this joint was previously related to the damping behavior of the leg during level locomotion ^3^, we can speculate, based on the PVPP model, that the combined action of the hip and the TMP joints might control gait-changes between grounded and aerial running as they regulate, respectively, the effective leg length and damping ratio during the stance.

To cope with drops of about 25% to 50% of leg length, the quail approached the perturbation more carefully and relied on double support. However, animals’ strategies to negotiate drops of 25% and 50% leg length differed. When negotiating visible drops of 25% leg length, the quail displayed rather subtle changes in the trailing leg, even though its effective length was longer than during level locomotion. This observation is supported by the slightly more extended hip and knee joints during stance, and a stiffer INT joint (less flexion-extension than level locomotion for assumed similar ground reaction forces), which might also have induced a vaulting descending motion of the CoM towards the lowered substrate.

To cope with visible drops of 50% leg length, the trailing leg displayed a more crouched configuration, and was less retracted than during level locomotion (Fig. 2). The shorter effective leg was produced by a significantly more flexed hip, INT and knee joints. Leg retraction displayed a trade-off between flexion of the hip, which protracted the leg, and of the knee, which in turn induced the contrary motion.

Thus, the quail used a large hip extension to extend the effective leg during stance but did not use a larger hip flexion to shorten it. This can be explained by the fact that hip extensor torque must be sufficient to stabilize a pronograde trunk and the overall locomotion ^18,35,42^.

At TD and during later stance, the trailing whole leg was nearly vertically oriented for 25% and 50% visible drops. Such a leg orientation may help to prevent a collapse of the leg. For the largest drop, the hip was significantly more abducted (see Fig. 3). The described leg placement permitted the pelvis to be rotated towards the trailing leg (yaw motion) and tilted towards the leading leg (roll motion) while descending towards the lowered substrate (Fig. 4).

#### Leading limb (stride i)

The leading effective leg touched down significantly later when stepping down, if compared to the same event during level locomotion. The angle of attack (α_0_) was steeper but did not vary with drop-height. At the same time the retraction of the trailing limb in the late stance was step-height related. This indicates that leg retraction velocity was decoupled from the trailing leg after crossing to the ground level, as observed in the aperture angle (see Fig. 2). This result suggests that the angle of attack and not the aperture angle is a target control parameter for leg placement when negotiating visible drops. During 1cm drops, the effective leg lengthening during swing is explained by hip extension, but especially by the significant extension of the TMP joint before TD. This shaped the subsequent behavior of the leg during stance. We think, that the more extended TMP joint at TD shifted spring-like behavior from the INT to the TMP joint (see Fig. 3). Gordon and colleagues showed that the guinea fowl displayed significantly higher activation of the *M. flexor perforatus digiti* III before and after their leg touched down in a sunken substrate ^9^. We speculate, that by preloading the tendons spanning the TMP joint during swing, the quail changed the viscoelastic properties of the joint (i.e., they shifted from a more damped joint behavior dominated by muscle properties to a more spring-like behavior dominated by elastic tissues, as observed in running humans ^43^ and turkeys ^44^. The goal of this anticipation seems to be two-fold. First, to maintain minimization of joint work under larger GRF and second, to reduce injury risk in soft tissues. By the way, this reflects the same strategy in experienced vs. unexperienced dogs in agility ^45^. The strategy of minimizing the sum of joint work also accounted for segmental bird kinematics in level locomotion ^46^. Recalling that joint work is the joint torque (T) times angular excursion, it follows that if the GRF increase, larger joint movements must be shifted to the joints located closer to the line of action of the GRF (having less torque). In addition, by shifting the spring-like behavior to the joint with a more convenient mechanical advantage ^47^, the quail may prevent soft tissue injuries by decreasing the tension in the tendons.

As was observed for drops of 10% leg length, the quail used a more extended leading leg (stride i) to negotiate drops of 25% leg length compared to level or 5 cm drops (see Table 2). However, the source of the leading leg lengthening was different from those depicted for drops of 10% leg length. The quail extended the INT joint instead of the TMP joint during swing (see Fig. 3). This simple change effected a dampened leg response after the drop. Focusing on the joint level, the TMP joint abandoned the spring-like behavior during stance depicted during 10% drops, and exhibited the dampened pattern described for level locomotion ^3^. It seems that the extension of the INT joint during swing permits muscular work to control leg compression and thus the energy dissipation after a visible drop. EMG data from the guinea fowl negotiating slow drops showed that the *M. gastrocnemius pars lateralis* was recruited earlier than the *M. flexores perforate digiti* III. This shift in the activation vanished for faster drops and level locomotion ^9^. Perhaps the onset in the activation of these muscles is used by birds to shape the viscoelastic response of the leg.

To negotiate 50% leg length drops, the aperture angle between the effective legs was similar to 25% leg length drops until the level line. However, after the leg crossed the level height, it was extended until TD. This indicates that the trailing limb rotated faster than the leading limb. Note that the slope of the mean leg angle before TD was quite flat until the level line (Fig. 2). Consequently, the retraction speed of the leading leg might be only slightly adapted when level TD is lost. At TD, the leading effective leg was shorter than in other drop conditions. Distal joint angles during 50% leg length drops were not significantly different from those exhibited by 2.5 cm drops. During this rather cautious drop negotiating technique, leg shortening seems to be performed by a more flexed hip joint at TD. During stance, the INT displayed a more bouncing-like behavior.

With increased drop height, the whole leg was more vertically oriented in the frontal plane and less abducted in the lowered substrate compared to unrestricted locomotion. This leg placement strategy prevented leg collapse and might have permitted the reorientation of the pelvis and thus the trunk in motion’s direction.

## Conclusions

To negotiate visible vertical perturbations, the quail reconfigured leg and joint kinematics related to perturbation type and height via different anticipatory strategies during swing and/or reactive control after TD. However, dramatic changes were observed only in the trailing limb for step perturbations of 50% of leg length. Leg and joint adaptations permitted the quail to regain steady-state locomotion already after one or two steps.

When coping with vertical perturbations, the quail adapted the trailing limb to permit that the leading leg steps on the elevated substrate in the same way as it does during level locomotion. This strategy may have reduced the need of reactive (feedback) response to readapt posture during leading leg’s stance.

The quail kept the function of the distal joints to a large extent unchanged during uneven locomotion, and most changes were accomplished in proximal joints. Up to middle step heights, hip extension was mainly used to lengthen the leg, or in combination with a more spring-like TMP joint to change to aerial running. However, to negotiate the largest visible step perturbations, all joints contributed to leg lengthening/ shortening in the trailing leg and both the trailing and leading legs stepped more vertically and less abducted. This indicates a sudden change in leg motor-control program. Further analysis is certainly necessary to understand muscle synergies, and overall neuromechanics underlining changes between dynamical and more safely gait programs.

## Methods

### Animals

Nine adult common quails [Phasianidae: *Coturnix coturnix* (Linnaeus 1758)] displaying a body weight ranging from 270 to 360 g were used for our analysis (see Table 11). The birds were housed at the Institute of Zoology and Evolutionary Research in Jena with access to food and water ad libitum. Housing, care, and all experimental procedures were approved by the Committee for Animal Research of the State of Thuringia (registry number 02-47/10). Animal keeping and experiments were performed in strictly accordance with the approved guidelines.

**Table 11.** Animals and strides

| Individual | Weight [g] | Strides |  |  |  |  |  |
| --- | --- | --- | --- | --- | --- | --- | --- |
|  |  | 1cm up | 2.5 cm up | 5 cm up | 1cm down | 2.5 cm down | 5 cm down |
| Schwarz | 341 |  | 1 | 5 | 1 | 5 | 2 |
| Rot | 284 |  | 3 | 4 |  | 4 | 1 |
| Silber | 295 | 1 | 5 | 2 | 2 | 2 |  |
| Dunkelgrün | 337 | 4 | 2 | 3 | 2 | 1 | 3 |
| Hellgrün | 277 |  | 3 |  |  |  |  |
| Lila | 362 | 1 |  |  |  |  |  |
| Rosa | 342 |  |  |  |  |  |  |
| Orange | 295 |  |  | 2 |  |  |  |
| Gelb | 307 | 3 | 2 |  |  | 4 | 3 |

### Experiments

For information about level locomotion experiments please refer to ^3^. In the step-up / step-down experiments, the quails moved across a 3 m long walking track at their preferred speeds. In the middle of the walking track, the birds negotiated visible drop/ step-up perturbations of 1.0 cm, 2.5 cm, and 5 cm. Those perturbations were created by supplementing the first (for drops) or the last (for step-up) half of the walking track. The track was covered with fine sheet rubber to reduce slipping. Body and limb kinematics were collected by using a biplanar X-ray fluoroscope (Neurostar, Siemens, Erlangen, Germany) at the facility of the Institute of Zoology and Evolutionary Research, Germany. X-ray sources were set to obtain recordings from the laterolateral and ventrodorsal projections. In addition, two synchronized standard light high-speed cameras (SpeedCam Visario g2, Weinberger, Erlangen, Germany) were used to cover both frontal and lateral perspectives of the track. The X-ray machine parameters were 40 kV and 53 mA, and a sampling frequency of 500 Hz. Raw video data was first undistorted by using a freely available MATLAB (The MathWorks, Natick, MA, USA) routine (www.xromm.org) provided by Brown University (Providence, RI, USA). As a base for the Automatic Anatomical Landmark Localization using Deep Features (see below), manual digitization of the joints and other landmarks [following ^3^] was performed using SimiMotion software (SimiMotion Systems, Unterschleißheim, Germany) on no more than five randomly distributed frames per trial.

### Automatic Anatomical Landmark Localization in Multi-view Sequences using Deep Features

In the following, the automatic multi-view landmark localization technique of the locomotion sequence is described, which is originally published in ^48^. Our method utilizes multi-view deep concatenated feature representations of annotated input images to train individual linear regressors for each view-based correspondent landmark pair. Based on a small number of annotated correspondent images of a multi-view sequence, the individual trained regressors locate all landmarks of the entire sequence in each view. In figure 6 the whole method pipeline is visualized. Afterwards, the automatic localized 2D landmarks of the dorsoventral and lateral view are utilized to reconstruct 3D landmark coordinates.

**Figure 6:**
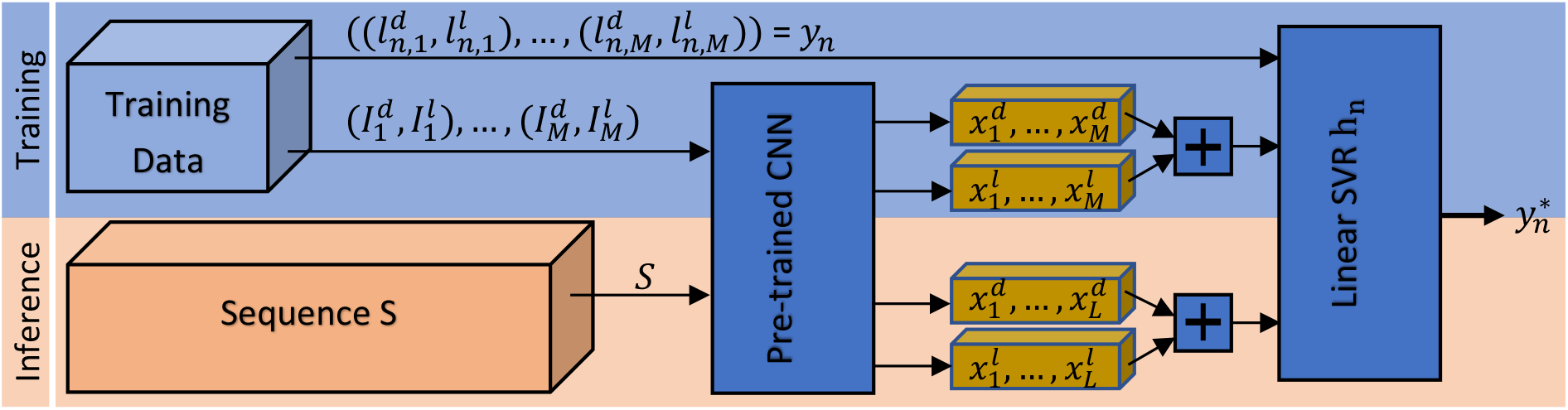
To train an individual multi-view landmark regressor *h_n_*, initially, the deep features 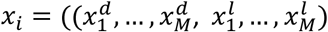 are extracted of *M* annotated image pairs. Afterwards, the concatenated features of correspondent image pairs serve as input for the regressor training. The landmark positions 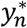 of unseen image pairs of *S* are predicted from the resulting trained model *h_n_*. This procedure is repeated for each of the *N* landmark pairs individually.

The utilized deep features are learned representations of images extracted from a Convolutional Neural Network (CNN) ^49^, which are mainly used for supervised computer vision tasks, like image classification, object recognition, or object tracking. The CNN learn in each of its convolutional layer several sets of individual convolutional filters based on the input images in the training process and provides thereby powerful feature representations of the utilized image domain.

The training of CNN models usually needs a lot of data, which is not available in our application. Hence, we choose a model of the AlexNet architecture ^50^ pre-trained on a similar task exploiting the same data domain of our application. This pre-trained model is trained for pose classification with the very same data of multi-view bipedal locomotion sequences to distinguish 10 quantized poses in each view during running on a trap. The semi-automatic annotation of the poses is described in ^48^. After training the CNN on the auxiliary task of pose classification, the CNN’s layer activations during inference can be exploited as deep features. In the following we describe the regressor training process for a single two-view locomotion sequence S utilizing the deep features.

The multi-view locomotion sequence *S* contains *L* correspondent image pairs from the dorsoventral and lateral view 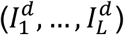 and 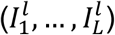. From each image pair 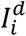 and 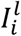 the deep features 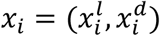 are extracted and concatenated from the fifth convolutional layer Conv-5 of the pre-trained CNN. Additionally, in *M* = 10 equidistant sampled frame pairs of both views, the correspondent *N* = 22 landmark position pairs *y* = (*y*_1_,…, *y_N_*) with 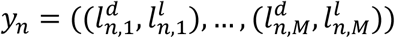 are annotated, which are used for single regressor training.

By utilizing each annotated corresponding landmark pairs *y_n_*, individual linear regressors *h_n_* are trained, which locates the correspondent landmarks in the remaining *L* – *M* images of both views, automatically.

As linear model *h_n_*, we train *N* single *ϵ*-SV regressors ^51^. Each linear regression model *h_n_* uses the given training data 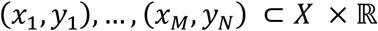 where *x_i_* denotes the deep features with 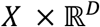 and *y_i_* the landmark positions of the *i^th^* landmark in the *M* frames. Hence, for each landmark position pair of both views, a single regressor hi is trained.

The goal of this regression task is to find a hyperplane *f*(*x*) = 〈*ω,x*〉 + *b* with a maximum deviation of *ϵ* from the target values *y_i_* for all training data. Given the fact that the vector *ω* is perpendicular to the hyperplane *f*(*x*), we only need to minimize the norm of *ω*, i.e., ║*ω*║^2^ = 〈*ω, ω*〉. When working with real data, in most cases, it is impossible to find a decent solution for this convex optimization problem based on potential outliers. With the addition of slack variables *ξ_i_* and 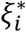 such infeasible conditions can be handled. We formulate the problem like ^51^:

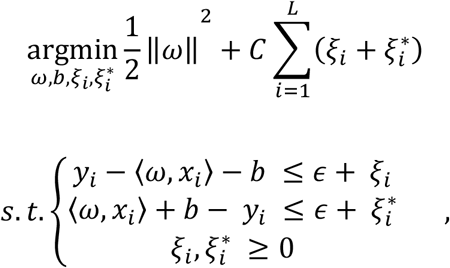

where *C* > 0 is a constant, which weights the tolerance of deviation greater than *ϵ*.

### C. Multi-view 3D Reconstruction

The dorsoventral and lateral 2-dimensional position data can be exploited to reconstruct these corresponded landmark points to 3-dimensional points in a metric space. To realize that a 3-dimensional calibration pattern in the form of a semi-transparent cube containing metal spheres is utilized, where each of the spheres have a distance of 1cm. By annotating at least seven individual corresponding spheres in both views, a relationship between the *annotated 2D pixel position* 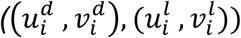 *to the 3D real word positions* (*X_i_, Y_i_, Z_i_*) *of* the spheres can be exploited. For more details on how *P* is estimated, we refer to ^52^.

#### Angle Calculation

Joint angles were computed as explained in ^3^, while model related leg kinematics following ^18,53^.

Three-dimensional kinematics (see Fig. 1 D): the pelvic local coordinate system was located in the centroid of the triangle composed by both hip joints and the pelvis cranial marker (*p_c_*). It measures the absolute motion of the pelvis related to the global coordinate system. It was defined by specifying first 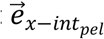 as an interim vector pointing from the right hip joint (*h_r_*) to the pelvis cranial marker 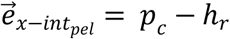, then 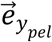 to be a vector pointing from *h_r_* to the left hip joint (*h_l_*), 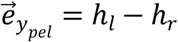, and 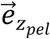 and 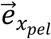 via cross-products as 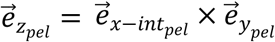 and 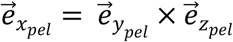. The whole-leg coordinate system measures the rotation of the whole leg related to the pelvis (estimates the three-dimensional rotations occurring at the hip joint). It was constructed as follows: 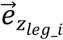 extends from the knee joint (*k_i_*) to the hip joint *h_i_* (right leg, _i = r_, left leg, _i=i_), e.g. 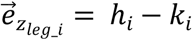. Then 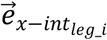 is an interim vector directed from TMP-distal markers (*tmp_dist_i_*) to *k_i_*, e.g., 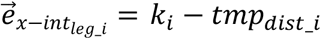. 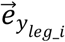 was then obtained as 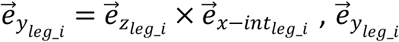 is hence perpendicular to the plane defined by the hip joint, the knee joint and the TMP-distal marker and points to the left (towards medial for the right leg and lateral for the left leg). Finally, 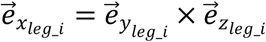. The whole-leg coordinate system was located in the middle of the femur (segment between hip and knee). To compute three-dimensional angles, we used the Cardan rotation sequence z-x-y. The left leg was used as reference. Thus, positive rotations around the x, y, and z axes represent, respectively, the inner rotation of the femur (whole leg rotates laterally), femoral retraction (hip extension), and femoral abduction. To build the mean using both legs, rotations around the z and the x axes for the right leg were multiplied by −1.

Kinematics were computed using a custom written script in Matlab 2017 (The MathWorks Inc., Natick, MA, USA).

#### Statistical analysis

Goal of our statistical analysis was to find kinematical differences effected by the different treatments. Following kinematic variables were defined as dependent variables: Global Parameters such as α_0_, ϕ_0_ and leg length, all joint angles and cardan angles for the pelvis and hip joint (relative angles between pelvis and leg). For the trailing limb, we analyzed the early stance (15%, because at TD in most of cases data was absent) and TO events. For the leading limb we analyzed the TD and the late stance (75%). In our analysis we included also the four precedents and the four following points relative to the selected event (event ± 4% of the stride).

Step locomotion are paired measures (same individuals) while step vs. level locomotion (grounded running) unpaired [level locomotion was collected in a different study, (Andrada et al., 2013b)]. For step locomotion repeated measures ANOVA was used to assess the influence of step-height and direction (up vs. drop) to the dependent variables. Post-Hoc tests with Bonferoni correction were afterwards performed to assess the influence of each treatment. Based on the homogeneity of the variances (Levene-test) we selected between TukeyHSD or Games-Howell tests. To test for significant differences between each step condition and level locomotion, we performed single multivariate ANOVAs (e.g., 2.5 cm step upwards vs. level).

Statistical analysis was implemented in R (Version: 3.5.3). We used the following libraries (R.matlab, data.table, stats, rstatix und car). To generate R-code we used the program “master” (free downloadable under https://starkrats.de).

## Declarations

### Ethics approval and consent to participate

All experiments were approved by and carried out in strict accordance with the German Animal Welfare guidelines of the states of Thuringia (TLV)

### Consent for publication

Not applicable

### Availability of data and materials

The datasets used and/or analyzed during the current study are available from the corresponding author on reasonable request.

### Authors’ contributions

E.A., M.S.F., and R.B. conceived the study. E.A and M.S.F supervised the experiments. J.D. and O.M. developed and O.M. performed the semi-automatic landmark identification, E.A. analyzed experimental data inclusive 2D and 3D kinematics, H.S. performed the statistics, E.A., M.S.F., D.J., M.T. and R.B. grants acquisition. E.A. drafted the manuscript. All authors contributed to the interpretation of the results and revised the manuscript.

## Acknowledgements

We would like to thank Lisa Dargel for animal training and animal guidance during the experiments. Rommy Petersohn and Yefta Sutedja for their technical assistance during the experiments. Ben Witt (formerly known as Ben Derwel) together with students worked hard to digitalize landmarks from the X-ray images for the semi-automatic identification.

